# *MKT1* alleles regulate stress responses through post-transcriptional modulation of Puf3 targets in budding yeast

**DOI:** 10.1101/2023.06.29.546914

**Authors:** Koppisetty Viswa Chaithanya, Himanshu Sinha

## Abstract

*MKT1* is a pleiotropic stress response gene identified by several quantitative trait studies with *MKT1*^89G^ as a causal variant, contributing to growth advantage in multiple stress environments. *MKT1* has been shown to regulate *HO* endonuclease post-transcriptionally via the Pbp1-Pab1 complex. RNA-binding protein Puf3 modulates a set of nuclear-encoded mitochondrial transcripts whose expression was found to be affected by *MKT1* alleles. This study attempts to relate the *MKT1* allele-derived growth advantage with the stability of Puf3 targets during stress and elucidate the roles of Pbp1 and Puf3 in this mechanism. Our results showed that the growth advantage of the *MKT1*^89G^ allele in cycloheximide and H O was *PBP1*-dependent, whereas in 4NQO, the growth advantage was dependent on both *PUF3* and *PBP1*. We compared the mRNA decay kinetics of a set of Puf3 targets in multiple stress environments to understand the allele-specific regulation by *MKT1*. In oxidative stress, the *MKT1*^89G^ allele modulated the differential expression of nuclear-encoded mitochondrial genes in a *PBP1* and *PUF3*-dependent manner. Additionally, *MKT1*^89G^ stabilised Puf3 targets, namely, *COX17*, *MRS1*, and *RDL2,* in an allele and stress-specific manner. Our results showed that COX17, MRS1, and RDL2 had a stress-specific response in stress environments, with the *MKT1*^89G^ allele contributing to better growth; this response was both *PBP1* and *PUF3*-dependent. Our results indicate that the common allele, *MKT1*^89G^, regulates stress responses by differentially stabilising Puf3-target mitochondrial genes, which allows for the strain’s better growth in stress environments.

**TAKE AWAY:** *MKT1* alleles vary stress responses by post-transcriptional modulation of Puf3 targets. *PBP1* and *PUF3* influence the regulation of Puf3 target stability under stress.

Nuclear-encoded mitochondrial Puf3 targets *COX17, MRS1* and *RDL2,* contribute to the growth advantage of *MKT1*^89G^ allele in oxidative stress.

## INTRODUCTION

The environment, with its dynamic compositional changes, poses a consistent threat to the cellular homeostatic balance (Simpson et al. 2012). Cells display an array of survival mechanisms to deal with these fluctuations by deploying environmental stress response (ESR), where the normal physiological course is transiently substituted with the synthesis of ESR-associated proteins (Lackner et al. 2012; Causton et al. 2001; Buchan and Parker 2009; Buchan et al. 2011; Loll-Krippleber and Brown 2017). Cellular regulation of gene expression during stress conditions predominantly employs post-transcriptional and translational controls to modulate their synthetic activity (Lackner et al. 2012; Martínez-Salas et al. 2013). RNA binding proteins associated with specific transcripts post-transcriptionally regulate their expression by modulating their localisation, degradation and translation (Saint-Georges et al. 2008; Martínez-Salas et al. 2013; Kechavarzi and Janga 2014; Klaus et al. 2014). Members of the Pumilio-Fem3 binding factor (PUF) family of RNA binding proteins exhibit this type of post-transcriptional regulation of sets of transcripts specific for each of five Puf protein subtypes (Olivas and Parker 2000; García-Rodríguez et al. 2007; Wang et al. 2018). Puf3 targets a module of about 220 transcripts that majorly includes nuclear-encoded mitochondrial proteins, which differ in their expression patterns during stress conditions influencing mitochondrial biogenesis (Gerber et al. 2004; Saint-Georges et al. 2008). While utilising carbon sources that demand active respiration, the repressor-like activity of Puf3 binding to the 3’UTR elements on the transcript decreases their expression by promoting degradation or preventing it (Olivas and Parker 2000; Miller et al. 2014).

Various stress conditions, including osmotic, oxidative, nutritional deprivation, chemical, high temperature, etc., were used to understand ESR in *Saccharomyces cerevisiae* (Causton et al. 2001; Gasch and Werner-Washburne 2002; Morano et al. 2012). Studies in yeast strains from clinical and natural environments have linked genetic diversity to varying stress responses (Liti et al., 2009).

Mapping studies in segregating populations to determine causal loci for stress responses identified *MKT1*, a pleiotropic stress response gene mediating high-temperature growth (Steinmetz et al. 2002; Sinha et al. 2008), sporulation efficiency (Deutschbauer and Davis 2005), chemical stress (Ehrenreich et al. 2010) and high ethanol concentration (Swinnen et al. 2012) in different yeast strains. Besides being causal for stress responses, *MKT1* is non-essential and alters mitochondrial stability, affecting stress response (Wickner 1987; Dimitrov et al. 2009). At the molecular level, *MKT1* facilitates mating type switching in yeast mother cells by selective post-transcriptional regulation of *HO* endonuclease during budding. Mkt1 interaction with Pbp1, a protein binding to poly-A binding protein, Pab1, supports the role of Mkt1 in the post-transcriptional regulation of transcripts selectively based on the consensus sequence on 3’UTR of transcripts (Tadauchi et al. 2004). While S288c has *MKT1*^89A^, non-synonymous SNP change *MKT1*^A89G^ substituting D30G in the polypeptide is conserved among various natural isolates (Liti et al. 2009). Allele replacement studies between SK1 and S288c have shown *MKT1*(D30G) increases sporulation efficiency (Deutschbauer and Davis 2005). Furthermore, genome-wide RNA expression analysis hypothesised that *MKT1*^A89G^ SNP variation could be causal in altering the transcript stability of Puf3 module genes under stress conditions (Lee et al. 2009; Sun et al. 2016). Similarly, QTL coding for *IRA2* was linked to regulating Puf4 activity and the causal polymorphisms were known to affect the transcripts encoding nucleolar ribosomal RNA-processing factors (Smith and Kruglyak 2008).

While *MKT1* as a QTL influence the Puf3 targets, another locus, Pop7, positively regulates Puf3 activity (Fazlollahi et al. 2014). Additionally, overlapping targets between the Puf family proteins and the mechanism of network rewiring in the absence of Puf3 influence the expression of specific deletion phenotypes (Lapointe et al. 2017). In the current study, we link the allelic effects of *MKT1* with the stress-specific stability of Puf3 targets and elucidate the roles of *PBP1* and *PUF3* in this mechanism.

## MATERIALS AND METHODS

### Yeast strains and growth conditions

All the yeast strains used in the experiments are derivatives of the *Saccharomyces cerevisiae* S288c strain. The parental strains used were the ‘S’ strain (S288c-*MKT1*^89A^) and the ‘M’ strain (S288c-*MKT1*^89G^, Gupta et al. 2015). These strains were derived from the original strains obtained from Deutschbauer and Davis (2005). Briefly, the original strains were sequenced to confirm the allele replacements and then were backcrossed to the S288c strain for three generations to remove other mutations found in the strains. Gupta et al. (2015) used these backcrossed S and M strains for their whole genome gene expression analysis.

Strains were grown in standard YPD (1% yeast extract, 2% peptone, 2% dextrose) medium at 30°C. Plasmid-based drug cassettes amplified using pFa6A (kanMX4), pAG25 (natMx4), and pAG28 (hphMX4) as templates (Goldstein and McCusker 1999) were transformed to generate gene deletions in parental strains (Gietz et al. 1995). The stocks of H_2_O_2_ (H_2_O_2_, TCI Cat#H1222) and Cycloheximide (CYC, Sigma Cat#18079) were prepared in the water while FCCP (FCCP, TCI Cat#C3463) and 4-Nitroquinoline 1-oxide (4NQO, Sigma Cat#N8141) stocks were made in DMSO. These chemicals were added to YPD and used for stress conditions. The final concentrations used for each assay are given below.

### Spot dilution assay

A saturated culture from a single colony was obtained for 5ml of YPD after overnight incubation at 30°C. A dilution of 1:100 of saturated culture was used for cell counting using a haemocytometer. A serial dilution series ranging from 10^8^ to 10^3^ cells/ml was made for each strain. Five microliters of each dilution in the series were used to spot on YPD (control) and plates containing YPD along with stress agents (8% ethanol, 0.002% H_2_O_2_, 0.4µg/ml 4NQO), 0.25µg/ml CYC and 6µg/ml FCCP). The plates were incubated for four days at 30°C, following which the growth was recorded. Stress resistance was measured by comparing the number of spots with growth. All strains used for this study were diploid except for growth kinetics experiments, which were haploid. All the strains used in this study are given in Supplementary Table 1, and the primers in Supplementary Table 2.

### Growth kinetics

Turbid culture of strains in YPD was obtained in 96-well cell culture plates after 24h incubation at 30°C with 250rpm. Experiments were performed in 96-well cell culture plates, with each sample represented in triplicates of 200µl per well. Each strain was grown in YPD as a control for assays indicated with different chemicals CYC (0.1µg/ml), 4NQO (0.3µg/ml), H_2_ O_2_ (0.01%) and FCCP (1µg/ml). The plate was incubated for 42h at 30oC at 355cpm (orbital), and OD600 was recorded every 30 min using a BioTek EPOCH2 microplate reader. The growth curves were used to compute the growth rate and corresponding relative fitness. The relative fitness was defined as a ratio of the growth rate under the test condition to the growth rate in control (YPD) for the same strain.

### Fluorescence measurements

M and S strains with GFP-tagged *MKT1* were used for fluorescence studies to compare the levels of native protein expression. The strains with C-terminal GFP tagging were generated using pYM25 and selected with a Hygromycin marker (Janke et al. 2004). The overnight culture was used to inoculate fresh media and was grown till 1 OD absorbance at 600nm. The culture was spun down and washed three times with PBS (PH 7.4). The pellet was resuspended in 2ml fresh PBS, and the suspension was diluted to 1 OD. 300µl of cell suspension was pipetted in each well in a black opaque 96-well plate as well as 96-well transparent tissue culture plate to measure fluorescence at Em/Ex 485/510nm and absorbance at 600nm respectively using BioTek multimode plate reader (Synergy H1). PBS was used as blank, and a strain without GFP was used as a negative control. The fluorescence intensity of each well is divided by the corresponding absorbance to get normalised fluorescence intensity, which was represented as the average of biological replicates.

### RNA extractions

To study the temporal degradation kinetics of a candidate mRNA, 1,10-phenanthroline (Sigma Cat#131377) was used as a transcription inhibitor at a concentration of 200µg/ml in YPD and stress conditions. The overnight culture was measured at OD600 and inoculated to grow from 0.2-0.8 OD in YPD (control). For stress conditions, cultures were grown from 0.8-1.0 OD after adding stress-inducing agents to the required concentrations of CYC (3µg/ml), 4NQO (2µg/ml), H_2_O_2_ (0.15%), and FCCP (20µg/ml). Later, 1,10-phenanthroline at 200mg/ml concentration was added to control and stress cultures. Samples were collected every 15 min after administration of the drug, for which 10ml of culture corresponding to a particular time point was spun down at 8,000 rpm for 30 sec, and the pellet was snap-frozen using liquid nitrogen.

Frozen cell pellets transferred to 1.5ml tubes were added with 50µl of Phenol-Chloroform-Isoamyl alcohol (25:24:1 ratio, PCI) (Sigma Cat#71617), 50µl of Lysis buffer (50mM Tris pH 7-7.4, 130mM NaCl, 5mM EDTA, 5% SDS) and glass beads (approximately 2/3 of the pellet volume) was added to the tube. The tubes were then vortexed at maximum speed for 20 min at 4°C, followed by centrifugation at 13,000 rpm for 15 min at 4°C. The aqueous layer was transferred to pre-cooled 0.5ml tubes, and an equal volume of PCI was added. After vigorous mixing, the mix was centrifuged at 13,000 rpm for 10 min at 4°C. This step was repeated. The aqueous layer was extracted, and an equal volume of Chloroform-Isoamyl alcohol (24:1 ratio) was added. After vigorous mixing, the mix was centrifuged at 13,000 rpm for 10 min at 4°C. The step was repeated. The aqueous layer was removed and transferred to pre-cooled Eppendorf tubes containing 1/20th volume of 3M sodium acetate and two volumes of absolute ethanol. Total RNA was precipitated by inverting tubes and incubating them at -20°C for 30 min, followed by centrifugation at 13,000 rpm for 15 min. The pellet was washed using 200µl of 80% ethanol and centrifuged for 2 min without disrupting the pellet. The supernatant was discarded, and the pellet was air-dried on ice for 30 min.

The pellet was resuspended in a buffer containing TURBO DNase (Invitrogen Cat# 8167) following the manufacturer’s instructions to remove genomic DNA contamination. The concentration of RNA in the resulting solution was measured using Nanodrop; a ratio of approximately 2.0 for 260/280 and 260/230 was considered suitable for further studies. One microgram of the sample was run on 2% agarose gel with ethidium bromide used to assess the integrity of the RNA sample.

### Quantitative gene expression studies

According to the manufacturer’s instructions, 3.125µg of RNA was used for cDNA synthesis using random hexamers (Invitrogen Cat#N8080127) for priming and enzyme Superscript III (Invitrogen Cat#18080093). From which 25ng of cDNA was used per qPCR reaction of 10µl using KAPA SYBR FAST (Sigma Cat# KK4618) master mix and primers. The reaction was carried out with Analytikjena qTOWER3 in a two-step cycle of denaturation at 95°C, annealing and elongation at 60°C for 35 cycles. The Ct values from the experiments were used to calculate the respective ΔCt (Costa et al. 2013) and ΔΔCt values, which were used for further analysis. ΔCt was calculated as Ct_(gene of interest)_ – Ct_(endogenous control gene)_, while ΔΔCt was calculated as ΔCt_(treated sample)_ – ΔCt_(untreated control sample)_. ΔΔCt were obtained using ΔCt of the initial time point as a control sample for all the time points in that series after adding 1,10-phenanthroline. ΔCt values of tested genes were used in expression and linear regression analyses between *MKT1* allelic backgrounds. Temporal ΔΔCt values were used in differential stability analysis using Chow’s test (Chow 1960).

## RESULTS

### *MKT1* alleles contribute differentially to growth phenotype during stress

The phenotyping using serial dilutions (Figure 1A) showed that while in control conditions (YPD), there was no difference between the M and S strains, under stress growth conditions – ethanol, H_2_O_2_, FCCP, 4NQO, and CYC, the M strain grew better than the S. A series of concentrations ranging from 2-10% ethanol, 0.005-0.04% H_2_O_2_, 0.2-0.8µg/ml CYC and 4NQO, and 0.1-10 µg/ml FCCP were tested for differences in growth phenotype. These phenotyping results were confirmed using haploids with growth kinetic assays where no difference was observed between the M and S strains in YPD; in stress, the M had better growth rate and relative fitness than the S strain (Figures 1B, 1C). In both the spot dilution and growth kinetics assays, the concentration of stress-inducing agents was optimised to provide observable phenotypic differences between M and S backgrounds. While spot dilution assays in media containing ethanol showed a prominent growth difference between M and S strains, it was omitted from further experimentation due to its volatility and prolonged doubling time induced by the lack of glucose.

**Figure 1:**
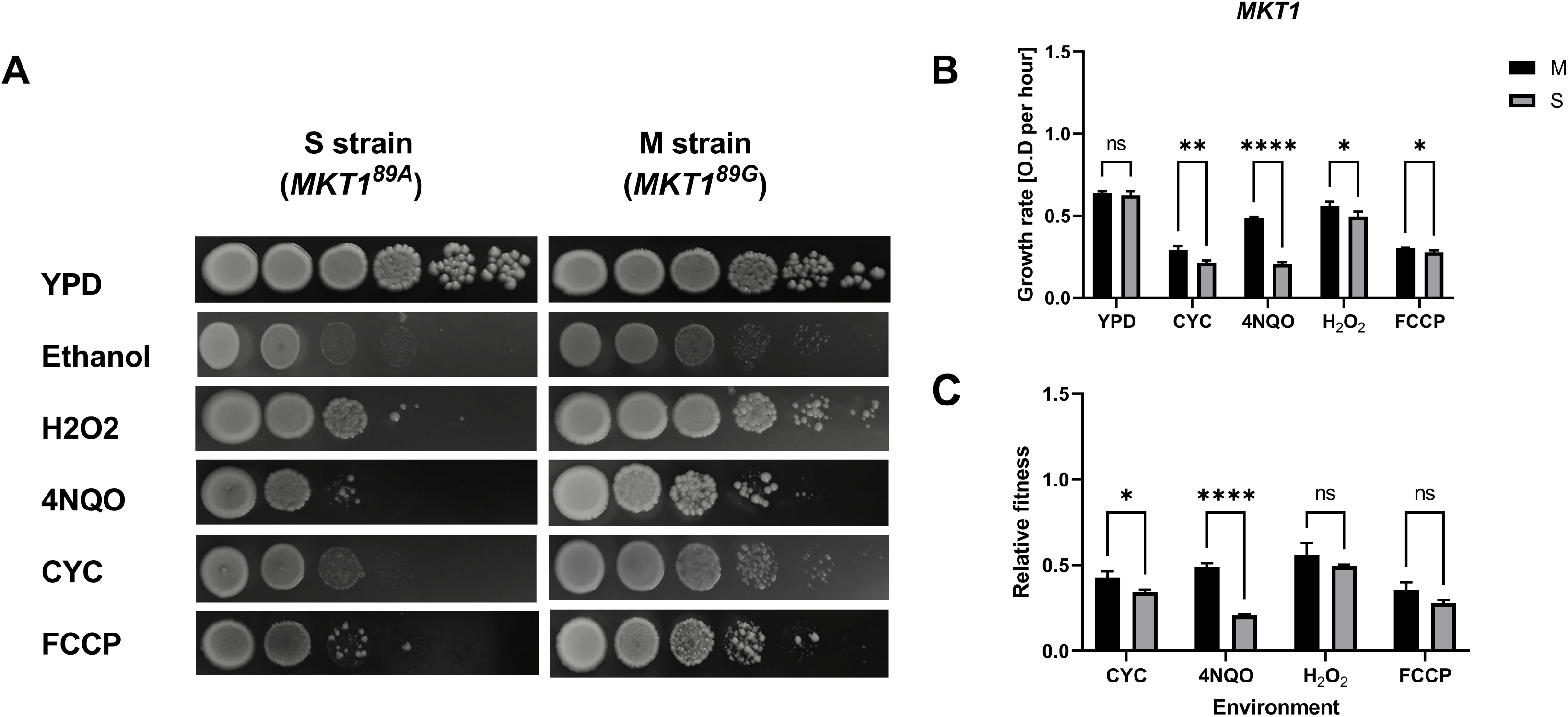
Analysis of *MKT1* allelic variants for pleiotropic stress resistance. [A] 10-fold serial dilution ranging from 10^8^-10^3^ cells/ml of S and M strains were spotted on YPD, 8% ethanol, 0.02% H_2_O_2_, 0.4µg/ml 4NQO, 0.25µg/ml CYC and 6µg/ml FCCP. Comparison of [B] growth rates and [C] relative fitness between M and S strains across YPD, 0.1µg/ml CYC, 0.3µg/ml 4NQO, 0.01% H_2_O_2_ and 1µg/ml FCCP. The experiments were performed in triplicates, and the error bars represent SD. P-values were calculated using a t-test, and significance was indicated as non-significant (ns), p < 0.05 (*), < 0.001 (**), < 0.0001 (***) and < 0.00001 (****) on the top of each comparison.

The phenotypic assays of the M and S strains showed that *MKT1* was dispensable in a stress-free environment, and the levels of *MKT1* in both backgrounds showed no difference (Figure S1, S2). However, the M strain was advantageous for growth rate and relative fitness in stress conditions over the S strain. While spot dilution experiments inferred a better phenotype of the M strain in H_2_O_2_ and FCCP, the relative fitness of both allelic strains was similar due to the lower chemical concentrations used for liquid cultures.

### *PBP1* and *PUF3* affect the magnitude and direction of *MKT1*^89G^ mediated growth advantage during stress

To understand the role of *PBP1* and *PUF3* in *MKT1*-mediated stress responses, *pbp1Δ*, *puf3Δ*, and *pbp1Δpuf3Δ* deletions were generated in M and S backgrounds. Growth kinetics assays in YPD showed that deleting *pbp1* or *puf3* or both had no effect in M and S backgrounds (Figure S3). Not surprisingly, the M-*pbp1Δ* strain had better relative fitness than S-*pbp1Δ* in CYC and 4NQO. However, the S-*pbp1Δ* strain showed better relative fitness than M-*pbp1Δ* in H_2_O_2_ and FCCP (Figure 2A).

**Figure 2:**
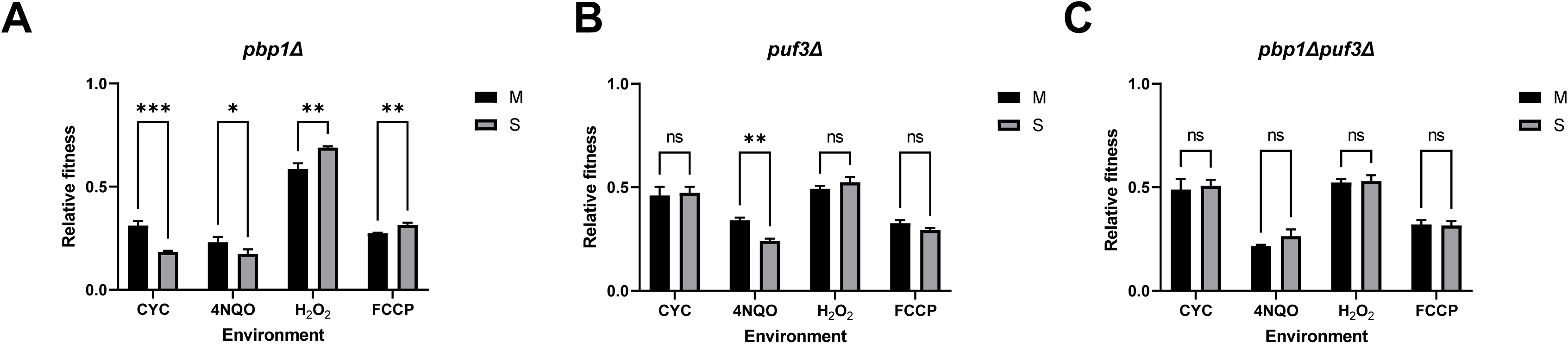
Role of *PBP1* and *PUF3* in *MKT1*-mediated stress responses. Relative fitness between M and S backgrounds with [A] *pbp1Δ*, [B] *puf3Δ* and [C] *pbp1Δpuf3Δ* deletions across YPD, 0.1µg/ml CYC, 0.3µg/ml 4NQO, 0.01% H_2_O_2_ and 1µg/ml FCCP. The experiments were performed in triplicates, and the error bars represent SD. P-values were calculated using a t-test, and significance was indicated as non-significant (ns), p < 0.05(*), < 0.001 (**), < 0.0001 (***), and < 0.00001 (****) on the top of each comparison.

No difference in relative growth was observed between M-*puf3Δ* and S-*puf3Δ* strains in YPD, CYC, H_2_O_2_, and FCCP (Figure S4). However, M-*puf3Δ* had higher relative fitness than S-*puf3Δ* in 4NQO (Figure 2B, Figure S2B).

As expected, double deletion strains of *pbp1Δpuf3Δ* in M or S backgrounds showed no growth difference in all the environments tested (Figure 2C). Thus, the double deletion of *pbp1* and *puf3* affected M and S backgrounds similarly, masking any individual allelic effects observed in the stress environments.

Analysing the *MKT1* allele and gene interactions across the environments showed that *PBP1* affected growth in CYC and H_2_O_2_, while *PBP1* with *PUF3* contributed to growth in 4NQO (Figure S4). This indicated that *MKT1-*mediated stress responses could employ both *PUF3* and *PBP1* in an environment-dependent manner.

### *MKT1* alleles control the expression levels of Puf3 targets using post-transcriptional machinery

We wanted to determine the expression levels of Puf3-target and Puf3-independent genes in each environment and background (M and S, Table 1, Supplementary Table 3). These genes were chosen to encompass diverse functional ranges like tRNA synthesis, splicing, translation and ribosomal proteins to find their involvement in different stress conditions. Therefore, we analysed the expression differences using ΔCt values of these genes at the initial time point between M and S backgrounds across multiple environments (Figure 3A).

**Figure 3:**
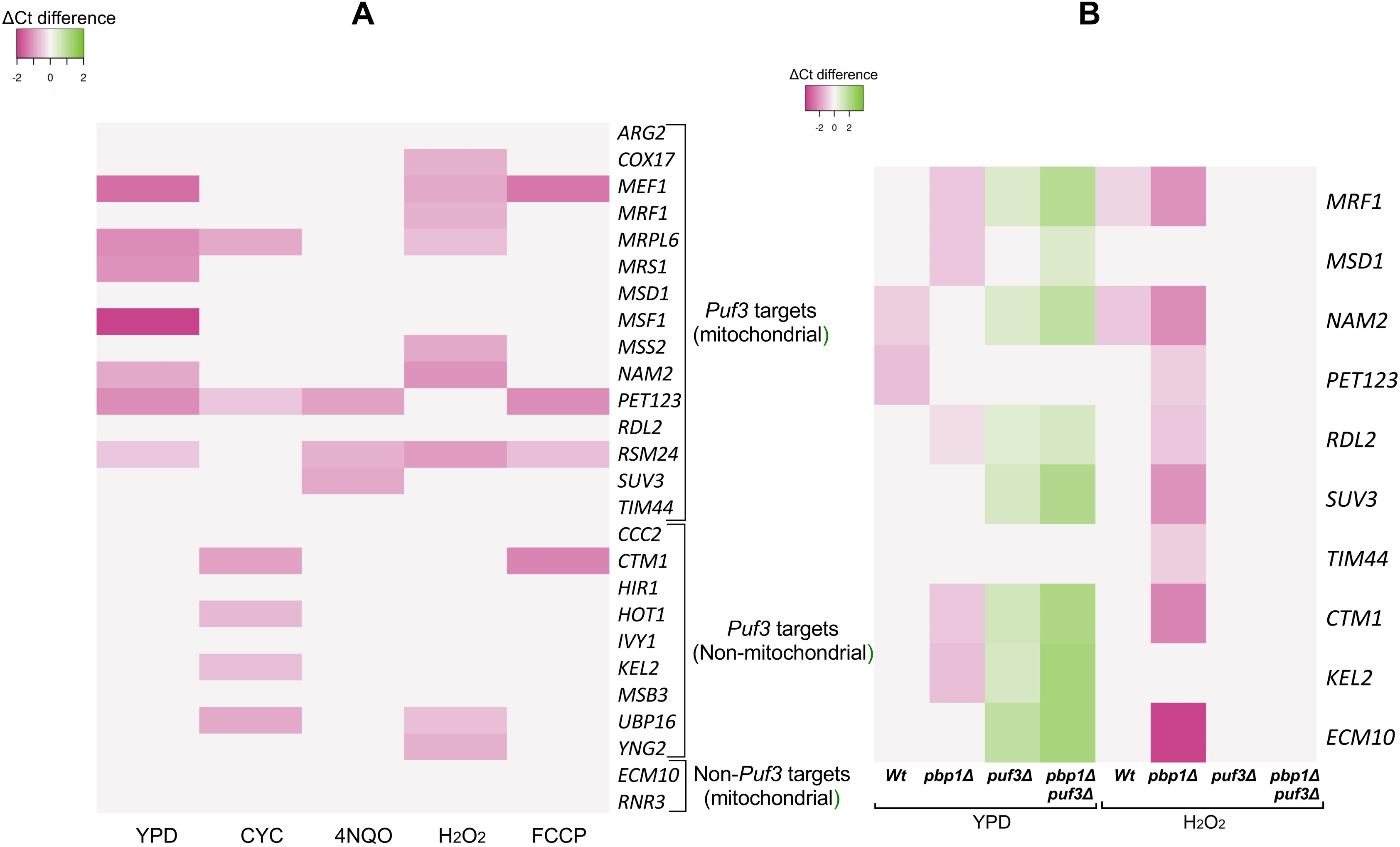
Differential expression (ΔCt) of Puf3 targets between M and S backgrounds at initial time point [A] wildtype across multiple environments, [B] *pbp1Δ*, *puf3Δ* and *pbp1Δpuf3Δ* deletions across YPD and H_2_O_2_. Cells were grown in YPD (control), CYC (3µg/ml), 4NQO (2µg/ml), H_2_O_2_ (0.15%) and FCCP (20µg/ml). The experiments were performed in triplicates, and the significant differences (p < 0.05) in ΔCt between M and S backgrounds were reported.

**Table 1:** List of genes analysed using RT-qPCR studies. Candidate Puf3 targets were selected from Gerber et al. (2004), and non-Puf3 targets were selected from SGD (. https://www.yeastgenome.org/).

| Puf3 affinity | Function | Gene |
| --- | --- | --- |
| Puf3 targets | Mitochondrial | *ARG2, COX17, MEF1, MRF1, MRPL6, MRS1, MSD1, MSF1, MSS2, NAM2, PET123, RDL2, RSM24, SUV3, TIM44* |
|  | Non-mitochondrial | *CCC2, CTM1, HIR1, HOT1, IVY1, KEL2, MSB3, UBP16, YNG2* |
| Non-Puf3 targets | Mitochondrial | *ECM10, RNR3* |

It was observed that some of the Puf3 targets, such as *MEF1, MRPL6, PET123,* and *RSM24*, showed higher expression in M compared to the S strain across three or more environmental conditions. In YPD, even though *MKT1* alleles did not show any growth differences, a few of the Puf3-target genes *MSF1, RSM24, NAM2, MRS1,* and *MEF1* were already differentially expressed at the initial time point. However, in CYC, 4NQO, H_2_O_2_, and FCCP, while there was a growth difference between M and S strains, some of the Puf3-targets genes were differentially expressed between the two strains, with M allele strains showing higher expression (Figure 3A). Furthermore, all genes with non-mitochondrial function had no expression difference between M and S strains in YPD (Table 2). The temporal ΔCt plots of each gene for all the conditions tested were given in Supplementary Figures S5-S9.

**Table 2:** List of genes showing differential expression of transcripts between M and S strains under each growth condition.

| Environment | Puf3 targets showing differential expression |
| --- | --- |
| YPD | *MEF1, MRPL6, MRS1, MSF1, NAM2, PET123, RSM24* |
| CYC | *MRPL6, PET123, CTM1, HOT1, KEL2, UBP16* |
| 4NQO | *PET123, RSM24, SUV3* |
| H<sub>2</sub>O<sub>2</sub> | *COX17, MEF1, MRF1, MRPL6, MSS2, NAM2, RSM24, UBP16, YNG2* |
| FCCP | *MEF1, PET123, RSM24, CTM1* |

In CYC and H_2_O_2_, where growth differences were observed between *MKT1* alleles, more genes were differentially expressed compared to other stress conditions. From the tested environments, including YPD, the expression levels in all the genes that showed differential expression were higher in the M compared to the S strain. This indicated that the M allele contributed to the differential expression by overexpressing some Puf3 targets in all the environments.

Interestingly, we observed that a few of the Puf3-target mitochondrial ribosome protein genes, *MRPL6, PET123* and *RSM24,* showed differential expression and stability in stress environments (Figure 3A, 3B). Several reports (Genuth and Barna 2012) indicate that ribosomal protein genes differentially translate a set of transcripts in a stress-dependent manner. Therefore, we did not analyse these genes further to avoid this ribosomal heterogeneity as a confounding factor.

It has been previously shown that *MKT1* activity depends on Pbp1 (Tadauchi et al. 2004). Furthermore, Lee et al. (2009) showed that *MKT1* played a role in the RNA stability of Puf3-dependent transcripts. To test if the differences described above depended on Puf3 and Pbp1, we deleted these genes in M and S backgrounds and measured expression levels at the initial time for a subset of Puf3-target genes (Figure 3B). This set of genes was selected based on the variable expression differences observed in wildtype in different growth conditions (Table 2). To obtain an unbiased estimate of the phenomenon due to the deletion of Pbp1 and Puf3, the established Puf3 target *COX17* (Olivas and Parker 2000) was omitted from this list of tested genes. As H_2_O_2_ showed the maximum number of differentially expressed genes relative to other stress conditions, we chose to study the roles of *PBP1* and *PUF3* in the H_2_O_2_ environment to compare against YPD control. In the *pbp1Δ* background, both in YPD and H_2_O_2_, for all the differentially expressed genes, the expression in M-*pbp1Δ* was higher than S-*pbp1Δ*. However, in the *puf3Δ* background, differentially expressed genes were observed in YPD alone, but interestingly, the expression of these genes was higher in the S-*puf3Δ* than in the M-*puf3Δ* strain. This is similar to the previous observations where Puf3 facilitated degradation in YPD but not under stress conditions (Miller et al. 2014). Comparison of the results between single deletions, *pbp1Δ* and puf3Δ, and double deletion *pbp1Δpuf3Δ* backgrounds showed that the differential expression of the genes was Puf3-dependent.

To understand if this differential expression of Puf3-target genes affected their transcript stability, the transcript levels were measured at regular intervals (15 min each till 60 min endpoint) after adding 1,10-phenanthroline. The ΔCt slopes of temporal samples of each Puf3-target gene were calculated using linear regression in the M and S strains and were compared in different environments. This comparison was made to determine if there was a change in the stability of the transcripts of Puf3-target genes for both wild-type M and S strains in YPD and H O , Puf3-target gene slopes were highly correlated (R^2^ = 0.83, p-value = 0.005) and (R^2^ = 0.78, p-value = 0.013), respectively). This observation suggested that in wild-type strains, all the Puf3-target gene transcripts behaved similarly for both environments, i.e., stabilised or degraded. For the *pbp1Δ* strain, in M and S backgrounds, Puf3-target gene slopes were only correlated in YPD (R^2^ = 0.66, p-value = 0.03) and not H_2_O_2_ (R^2^ = 0.18, p-value = 0.6). However, for the *puf3Δ* strain, in M and S backgrounds, only in H O , Puf3-target gene slopes were correlated (R^2^ = 0.89, p-value = 0.0005), whereas in YPD (R^2^ = 0.05, p-value = 0.87) no correlation was observed. These results indicated that in H O , Puf3-target gene transcripts were under Pbp1 regulation. The temporal ΔCt plots of each of the genes analysed under the deletion backgrounds of *pbp1Δ*, *puf3Δ*, and *pbp1Δpuf3Δ* for all the conditions tested were given in Supplementary Figures S10-S12.

### *PBP1* and *PUF3* interact differentially with *MKT1*^89G^ allele to stabilise Puf3 targets

Therefore, to check if these accessory proteins differentially altered transcript stability in an *MKT1* allele-specific manner in H_2_O_2_, we studied *pbp1Δ, puf3Δ and pbp1Δpuf3Δ* in the M and S backgrounds and compared transcript degradation patterns with wildtype using temporal RT-qPCR assay (Figure 4). Using the initial point sample as a control for the temporal qPCR data ΔΔCt values were computed, and their median was used as a metric to measure the stability of Puf3 targets after the transcriptional stop. In the wild type for YPD (p-value = 0.99) and H_2_O_2_ (p-value = 0.23), there was no significant difference between the median ΔΔCt of the M and S strains. However, when the median ΔΔCt values of the M strain were compared between YPD and H_2_O_2_, it was observed that the Puf3 targets were preferentially stabilised in H_2_O_2_ (p-value < 10^-3^, Figure 4A). This stabilisation of Puf3 targets was also observed in the S strain between YPD and H_2_O_2_ (p-value < 10^-3^, Figure 4A). This indicated that the degradation rates of Puf3 targets in M and S strains were similar in YPD and H_2_O_2_. However, in YPD, between M-*pbp1Δ* and S-*pbp1Δ* strains, there was a significant difference in the degradation rates for most Puf3 gene transcripts (p-value < 10^-3^, Figure 4B). The lower the median ΔΔCt value, the more stabilised the Puf3 transcripts. In H_2_O_2_, similarly, S-*pbp1Δ* had more stabilised transcripts than the M-*pbp1Δ* (p-value < 10^-3^, Figure 4B). However, when a comparison was made between M-*pbp1Δ* across YPD and H_2_O_2_, there was a significant difference in stability, with the transcripts in YPD being more stable than in H_2_O_2_ (p-value = 0.004, Figure 4B). This was not the case for transcripts across the two environments in S-*pbp1Δ* (p-value = 0.24, Figure 4B). This observation indicated that the role of Pbp1 in transcript stabilisation was more prominent in the M than in the S strain. In the *puf3Δ* deletion background, the observations were similar to the *pbp1Δ* background, with significant differences between the M and S backgrounds in YPD (p-value = 0.03) and H_2_O_2_ (p-value < 10^-3^). Puf3 is a known regulator of Puf3 targets in non-stress conditions like YPD (Olivas and Parker 2000). Therefore, we observed that in the absence of *puf3Δ* in the M background, the Puf3 transcripts were stabilised more than what was observed in the wildtype M background (Figure 4A and 4C). Finally, in the *pbp1Δpuf3Δ* deletion strains in the M and S backgrounds, the patterns of the Puf3 transcript stability remained similar to what was observed in the *puf3Δ* and *pbp1Δ* backgrounds. However, in this case, the absence of *pbp1Δpuf3Δ* in both backgrounds stabilised the Puf3 transcripts more than observed in the wildtype backgrounds (Figure 4A and 4D). Therefore, there was an environment-dependent stabilisation of Puf3-target genes in the M background, indicating that Pbp1 and *MKT1* alleles regulate mRNA levels post-transcriptionally.

**Figure 4:**
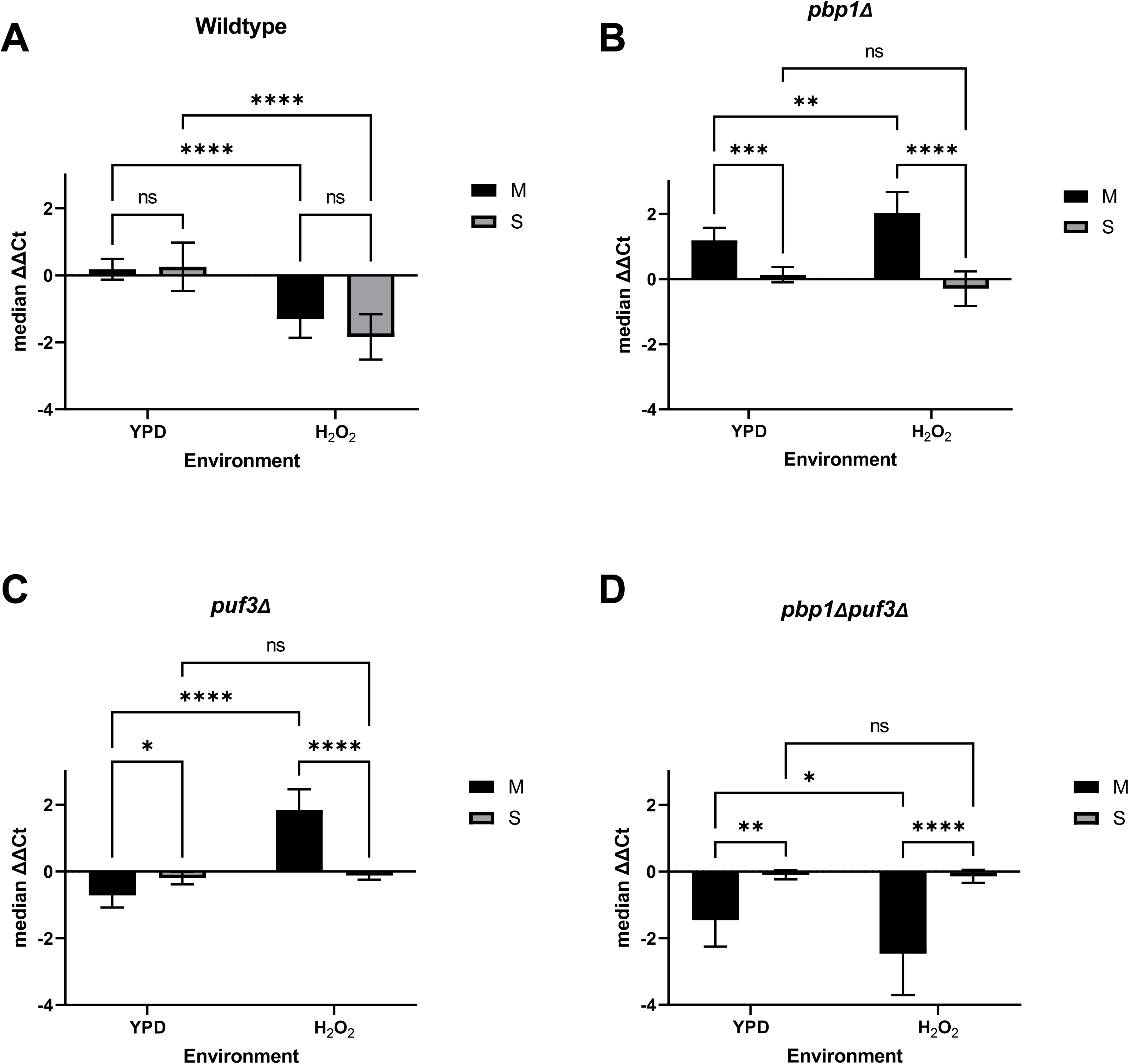
Analysis of temporal median ΔΔCt values among Puf3 targets between M and S backgrounds in [A] wildtype [B] *pbp1Δ*, [C] *puf3Δ* and [D] *pbp1Δpuf3Δ* deletions across YPD and H_2_O_2_ (0.15%). The experiments were performed in triplicates, and the error bars represent SD. P-values were calculated using ANOVA, and significance was indicated as non-significant (ns), p < 0.05 (*), 0.001 (**), 0.0001 (***) and 0.00001 (****) on the top of each comparison.

We wanted to test whether *MKT1* alleles differentially affect the stability of Puf3 targets across the environments. Temporal mRNA samples were analysed using transcript-specific RT-qPCR assay to monitor transcript decay rates for each transcript in the M and S backgrounds and across environments. The resulting temporal ΔΔCt values of each transcript in both M and S backgrounds were compared, and Chows’ test was used to determine significance. In YPD, all the genes except *HIR1, MSB3* and *MSD1* showed non-significant differences in the stability between the *MKT1* alleles. However, some Puf3-target gene transcripts showed significant variation in their stability rates between the *MKT1* alleles (Table 3). For example, in CYC, *COX17, MRS1, RDL2,* and 4NQO, *PET123, RDL2* , and *YNG2* significantly varied in their stability rates between the two *MKT1* alleles. The differentially stabilised transcripts belonged to mitochondrial and non-mitochondrial Puf3-target genes.

**Table 3:**
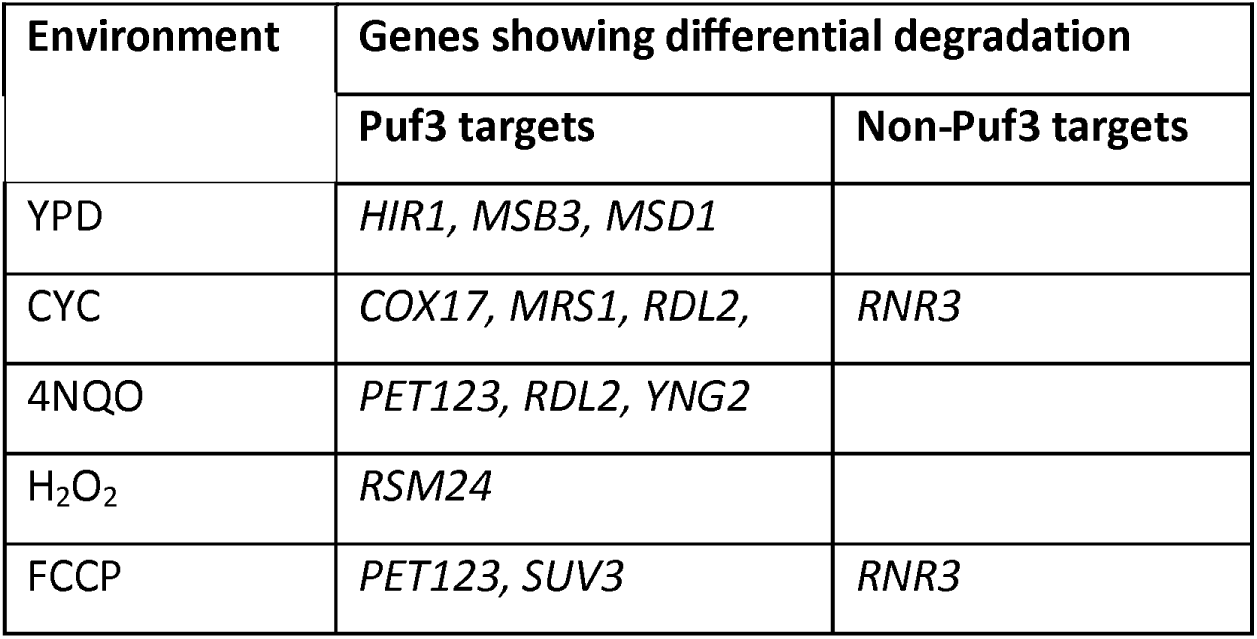
List of genes showing differential degradation of transcripts between M and S strains under each growth condition.

| Environment | Genes showing differential degradation |  |
| --- | --- | --- |
|  | Puf3 targets | Non-Puf3 targets |
| YPD | *HIR1, MSB3, MSD1* |  |
| CYC | *COX17, MRS1, RDL2,* | *RNR3* |
| 4NQO | *PET123, RDL2, YNG2* |  |
| H<sub>2</sub>O<sub>2</sub> | *RSM24* |  |
| FCCP | *PET123, SUV3* | *RNR3* |

### *COX17*, *MRS1* and*RDL2* are essential for the M strain to maintain growth advantage in stress environments

To understand the role of differentially expressed Puf3 targets in the pleiotropic stress response of MKT1 alleles, we curated a subset of Puf3 targets to study their roles by deleting them in both M and S backgrounds. Puf3 targets - *COX17, IVY1, MRF1, MRS1, MSD1,* and *RDL2* were selected based on their differential expression and varying stability between the *MKT1* alleles observed in qPCR assays. *COX17* and *MRS1* were observed to be both differentially expressed and differentially stabilised. *MRF1* represents differential expression alone. Similarly, *MSD1* and *RDL2* have shown differential degradation alone. *IVY1* was selected as a non-mitochondrial gene with no differential expression between *MKT1* alleles. *COX17* acts as a copper metallo-chaperon in mitochondria (Heaton et al. 2000). Phospholipid-binding protein *IVY1* regulates vacuole fission (Lazar et al. 2002). *MRF1* is a mitochondrial translation release factor (Pel et al. 1992). RNA-binding protein *MRS1* controls RNA processing via splicing Group I introns in mitochondria (Kreike et al. 1986). *MSD1* is a mitochondrial aspartyl-tRNA synthetase (Gampel and Tzagoloff 1989). *RDL2* is a mitochondrial thiosulfate sulfurtransferase (Foster et al. 2009).

We grew these gene deletion strains in CYC (0.1µg/ml), 4NQO (0.3µg/ml), and H_2_O_2_ (0.01%) and compared their growth between M and S across wildtype, *pbp1Δ*, *puf3Δ*, and *pbp1Δpuf3Δ* backgrounds for each environment. The concentrations of these chemicals in these liquid culture assays, even within the range tested, were lower than those used in the spot dilution assays. Lower concentration was used in liquid culture to allow these wildtype and deletions strains to grow in liquid media. The phenotype difference between M and S in wildtype was used to compare the effect of specific gene deletion in the respective environment.

In the wild type, as shown previously, the M strain grew better than the S in CYC (Figure 5A). When *PBP1* was deleted, the M strain was still better than the S. However, the fitness advantage for the M strain over the S was lost when either *puf3Δ* or *pbp1Δpuf3Δ* were deleted. This indicated that in CYC, the growth advantage of the M allele was *PUF3*-dependent, and *PUF3* was epistatic over *PBP1*. In 4NQO, in the wild type, the fitness advantage of the M strain over the S was independent of *PBP1* and *PUF3*, but when both these genes were deleted, no difference was observed (Figure 5E). This observation showed that in 4NQO, both *PBP1* and *PUF3* had an additive effect on growth. In the wildtype, no significant growth differences were observed in H_2_O_2_ for any strain.

**Figure 5:**
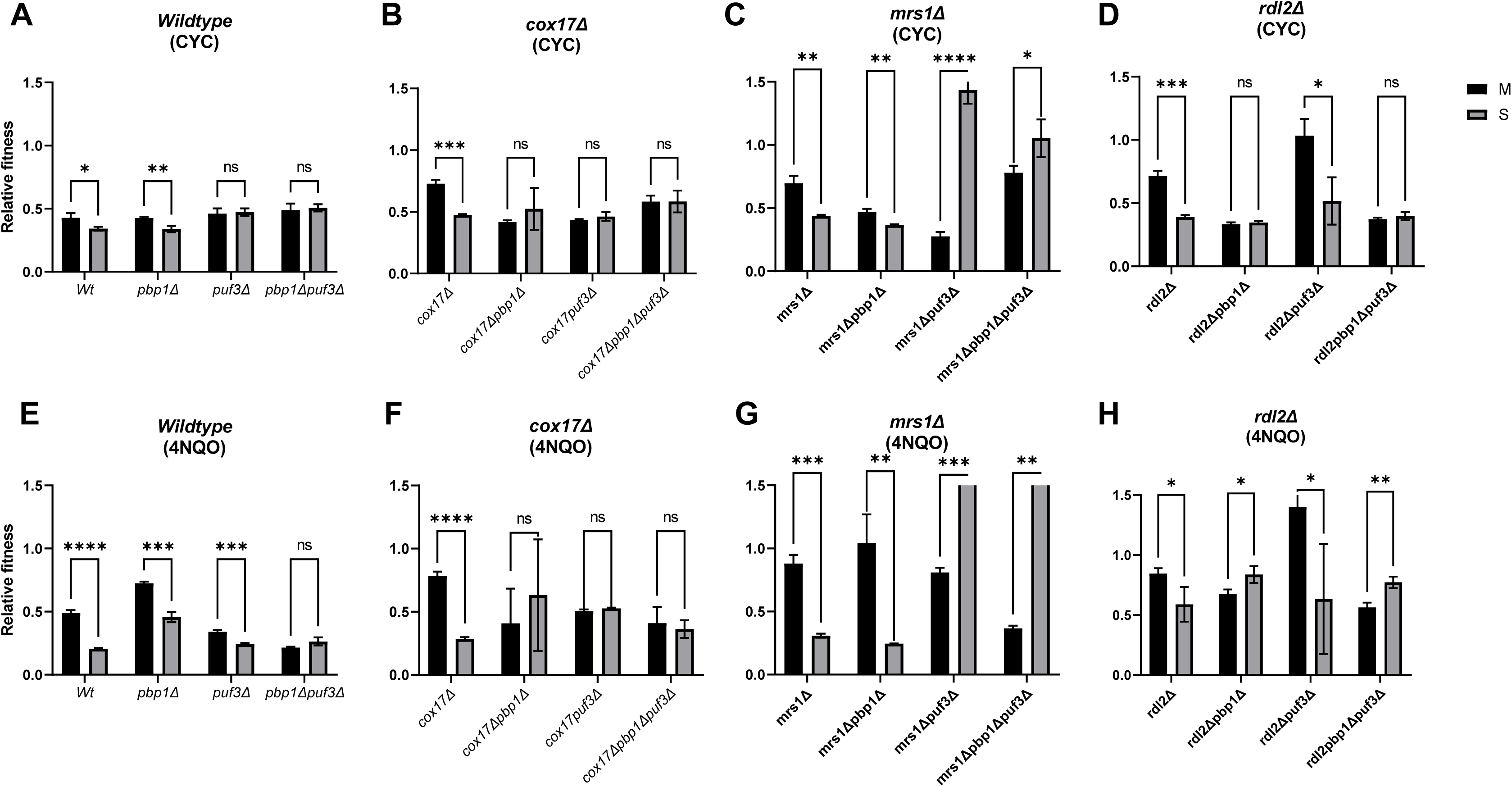
Effect of *COX17*, *MRS1* and *RDL2* on *MKT1* allelic response in CYC and 4NQO. Comparison of relative fitness between *MKT1* alleles in wildtype*, cox17Δ, mrs1Δ* and *rdl2Δ* backgrounds and the effect of wildtype, *pbp1Δ, puf3Δ* and *pbp1Δpuf3Δ* for each deletion in [A-D] 0.1µg/ml CYC and [E-H] 0.3µg/ml 4NQO. The experiments were performed in triplicates, and the error bars represent SD. P-values were calculated using a t-test, and significance was indicated as non-significant (ns), p < 0.05 (*), 0.001 (**), 0.0001 (***) and 0.00001 (****) on the top of each comparison.

In the *cox17Δ* background in both the M and S strains, the roles of *PBP1* and *PUF3* were altered. Compared to the wild type, *PBP1* and *PUF3* were required in CYC for the growth advantage of the M allele (Figure 5B). Therefore, *PUF3* was no longer epistatic over *PBP1* in this background. Similarly, in 4NQO, compared to the wild type, no growth difference was observed in either *pbp1Δ* or *puf3Δ* (Figure 5F). This indicated that *COX17* was required for *PBP1* and *PUF3*-dependent growth advantage of the M strain in all three environments.

In the *mrs1Δ* background, similar to the wild type, in CYC and 4NQO, the deletion of *pbp1Δ* resulted in a growth advantage of the M strain over the S (Figure 5C, G). However, in contrast to the wild type, the deletion of *puf3Δ* resulted in a reversal of the growth advantage, with the S strains growing better than the M. This reversed phenotype was also observed in the *pbp1Δpuf3Δ* double deletion background. This indicated that first, the growth advantage of the M allele was *PUF3*-dependent, and second, in the *mrs1Δ* background, *PUF3* was required for the activity of the M allele.

In the *rdl2Δ* background, in CYC, the growth advantage of the M strain was dependent on *PBP1*, with *PBP1* being epistatic over *PUF3* as the double deletion *pbp1Δpuf3Δ* was similar to *pbp1Δ* alone (Figure 5D). In 4NQO, again, the effect of *RDL2* was *PBP1*-dependent (Figure 5H). Interestingly, in 4NQO, deletion *pbp1Δ* resulted in the S strain having better growth than the M. Compared to the *MRS1* result, where better growth of the S strain was found to be dependent on *PUF3*, here in *RDL2*, it was dependent on *PBP1*. This indicated that in the same environment, the effects and roles of *PBP1* and *PUF3* are genetic background dependent.

In the H_2_O_2_ environment, there was no difference between the M and S strains in the wildtype, even when *pbp1Δ* and *puf3Δ* were deleted singly or in combination (Figure 6A). Interestingly, in *cox17Δ, mrs1Δ* and *rdl2Δ* backgrounds, the M strain was better than the S (Figure 6B, 6C, 6D). But, when in these backgrounds, either *pbp1Δ* or *puf3Δ* or *pbp1Δpuf3Δ* were deleted, the growth advantage of the M strain was lost. This indicated that the *Puf3*-dependent nuclear-encoded mitochondrial genes were required for better growth of the S strain, meaning that the non-functional allele of *MKT1* needed these genes for proper growth. This conclusion was further supported by deletion phenotypes of *msd1Δ* and *mrf1Δ,* which showed a similar phenotype pattern as *cox17Δ* (Figure S13F, 13I). For a non-mitochondrial gene, like *IVY1*, the deletion of *ivyΔ* was the same as wildtype phenotype, indicating that this phenotypic effect might be specific to *Puf3*-dependent mitochondrial genes only (Figure S13C).

**Figure 6:**
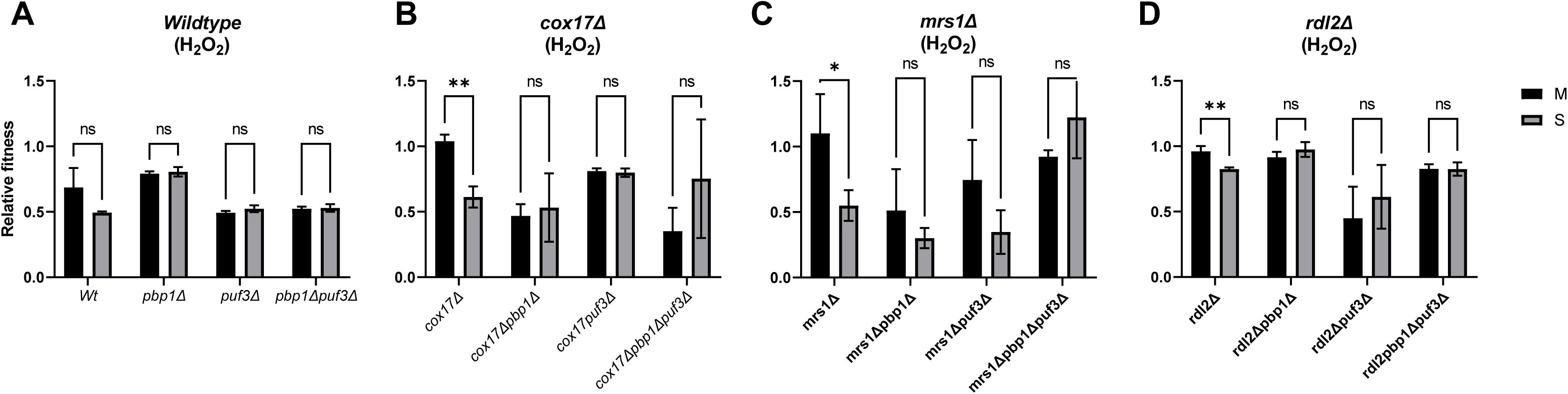
Effect of *COX17*, *MRS1* and *RDL2* on *MKT1* allelic response in H_2_O_2_. Comparison of relative fitness between *MKT1* alleles in wildtype*, cox17Δ, mrs1Δ* and *rdl2Δ* backgrounds and the effect of wildtype, *pbp1Δ, puf3Δ* and *pbp1Δpuf3Δ* for each deletion in [A-D] 0.01% H_2_O_2_. The experiments were performed in triplicates, and the error bars represent SD. P-values were calculated using a t-test, and significance was indicated as non-significant (ns), p < 0.05 (*), 0.001 (**), 0.0001 (***) and 0.00001 (****) on the top of each comparison.

## DISCUSSION

Our work was based on the hypothesis that *MKT1* allele-mediated stress resistance relied on post-transcriptional modulation of mitochondrial activity via Puf3 targets. While establishing the individual roles of *PBP1* and *PUF3* as interacting genes, we attempted to provide mechanistic insights for these allelic interactions. From the tested environments ranging from oxidative, genotoxic, translational stress, Cadmium chloride, and high-temperature growth, we identified the diverse stress conditions where the wildtype M strain consistently grew better than the S strain (Figure 1, S14).

Interestingly, Puf3 upregulated the expression of a few of its mitochondrial targets in stress conditions specific to the M strain, indicating the role of mitochondria in *MKT1* allele-specific stress resistance (Figure 4). A few of these mitochondrial target genes were observed to be crucial for the M strain exhibiting better fitness in diverse stress conditions. This difference in fitness was dependent on *PBP1* or *PUF3* or both in an environment-specific manner (Figure 5, 6). This indicated that the role of the *MKT1* allele on fitness was both dependent on the environment and on the presence of *PBP1* and *PUF3*.

The formation of the Mkt1-Pbp1-Pab1 complex facilitates post-transcriptional regulation by Mkt1. The growth phenotype of the S strain was different from S-*pbp1Δ* in CYC and H_2_O_2_, indicating the formation of the Mkt1-Pbp1-Pab1 complex remains unaffected between M and S alleles under stress (Figure S4). As Pbp1 localise to stress granules regulating transcript deadenylation (Swisher and Parker 2010), this particular association allows Mkt1 to control the stress-specific stability of mRNA. Mkt1, with its uncharacterised interaction with Puf3, was observed regulating Puf3 target degradation in an allele-specific manner. Puf3, while binding to its target mRNA, governed their access to translation machinery and was known to transiently repress nuclear-encoded mitochondrial transcripts in oxidative stress (Rowe et al. 2014). Our analysis with Puf3 target deletion strains in diverse environments confirmed that the allele-specific phenotype of *MKT1* was both Pbp1 and Puf3 dependent. This indicates that the ability of the M allele to stabilise stress-specific mitochondrial transcripts to modulate mitochondrial activity accounts for its better stress responses than the S allele. However, the nature of the interaction between the *MKT1*^89G^-Pbp1-Pab1 complex and Puf3 to specify the fate of a transcript is unknown. The possible mechanisms by which the Puf target transcript can be regulated include interactions from other Puf proteins exemplified in *ZEO1* mRNA interacting with both Puf1 and Puf2 (Haramati et al. 2017), facilitating mRNA degradation by promoting deadenylation (Olivas and Parker 2000) and mRNA localisation to mitochondria via Mdm12 and Tom20 (Miller et al. 2014).

Our growth experiments have established the qualitative effect of the M-allele’s growth advantage compared to the S-allele across multiple stress conditions in wildtype and most of the Puf3 target deletion backgrounds (Figure 5, 6). However, the magnitude of growth advantage, i.e., the difference between the fitness of M and S backgrounds, was variable with respect to each deletion across the environments (Figure S15). For example, the effect of deletion of these mitochondrial genes *COX17*, *MRS1* and *RDL2* was variable with respect to the environment and the allelic background. The influence of these genes on the growth difference between the M and the S strains was higher in H_2_O_2_ compared to CYC and 4NQO. One of the interesting observations was that *IVY1,* as a non-mitochondrial Puf3 target, had a phenotype similar to the wildtype in H_2_O_2_.

Our results highlight another level of regulatory complexity where coding polymorphisms in QTL, like *MKT1*, can modulate multiple stress responses through post-transcriptional control in an environment-specific manner.

## ACKNOWLEDGEMENTS

The authors acknowledge help from Rajeeva Lokshanan in the statistical analysis of some of the data. The authors thank K Subramanian for critically reading the manuscript and comments. VCK acknowledges the University Grant Commission Junior Research Fellowship. HS acknowledges intra-mural funding from IIT Madras and Excelra Knowledge Solutions Pvt. Ltd. (CR/22-23/0026/BT/EXCE/008752).

## AUTHOR CONTRIBUTIONS

VCK and HS designed all the experiments, VCK performed all the experiments, and VCK and HS wrote the manuscript.

## SUPPLEMENTARY INFORMATION

### Supplementary Tables

**Supplementary Table S1**: List of strains used in this study.

**Supplementary Table S2**: List of primer sequences used in this study.

**Supplementary Table S3** : List of Puf3 and non-Puf3 target genes used for RT-qPCR assays with their annotations.

**Supplementary Table S4**: The differential expression of ΔCt for wildtype in YPD, CYC, 4NQO, H_2_O_2_, FCCP, and for the deletion strains in YPD and H_2_O_2_.

### Supplementary Figures

**Figure S1**: Non-essentiality of *MKT1*. 10-fold serial dilution ranging from 10^8^-10^3^ cells/ml of M, S and S288c *mkt1Δ* strains were spotted on YPD.

**Figure S2** : Expression of *MKT1* at transcript and protein levels in M and S strains. [A] ΔCt expression levels of *MKT1* in M and S strains grown in YPD control and 0.15% H_2_O_2_. [B] Normalized fluorescence intensity of M (*MKT1*^89G^-GFP), S (*MKT1*^89A^-GFP) and Negative control (S strain) grown in YPD and H_2_O_2_. Strains with GFP-tagged *MKT1* were measured for fluorescence at Em/Ex 485/510 and normalised with their absorbance at 600 nm. Each experiment was performed with three biological replicates, and the resultant mean was plotted, and the error bars represent SD. P-values were calculated using a t-test, and significance was indicated as non-significant (ns), p < 0.05 (*), 0.001 (**), 0.0001 (***) and 0.00001 (****) on the top of each comparison.

**Figure S3**: Growth rate differences between M and S backgrounds in YPD across *Wt*, *pbp1Δ, puf3Δ* and *pbp1Δpuf3Δ.*The experiments were performed in triplicates, and the error bars represent SD. P-values were calculated using a t-test, and significance was indicated as non-significant (ns) at the top of each comparison.

**Figure S4** : Role of *PBP1* and *PUF3* on the fitness of M and S backgrounds in *Wt*, *pbp1Δ, puf3Δ* and *pbp1Δpuf3Δ.* Relative fitness between M and S backgrounds across [A] 0.1 µg/ml CYC, [B] 0.3 µg/ml 4NQO, [C] 0.01% H_2_O_2_ and [D] 1 µg/ml FCCP. The experiments were performed in triplicates, and the error bars represent SD. P-values were calculated using ANOVA, and significance was indicated as non-significant (ns), p < 0.05 (*), 0.001 (**), 0.0001 (***) and 0.00001 (****) on the top of each comparison.

**Figure S5**: Comparison of ΔCt values across five time points of multiple genes between M and S strains grown in YPD. T=0 is the time of addition of 1,10-phenanthroline. Time points differ in 15 min with the consecutive one. Each represents as the mean of three biological replicates, and the error bars represent SD.

**Figure S6**: Comparison of ΔCt values across five time points of multiple genes between M and S strains grown in 0.15% H_2_O_2_. T=0 is the time of addition of 1,10-phenanthroline. Time points differ in 15 min with the consecutive one. Each sample represents the mean of three biological replicates, and the error bars represent SD.

**Figure S7**: Comparison of ΔCt values across five time points of multiple genes between M and S strains grown under 2µg/ml 4NQO. T=0 is the time of addition of 1,10-phenanthroline. Time points differ in 15 min with the consecutive one. Each represents the mean of three biological replicates, and the error bars represent SD.

**Figure S8**: Comparison of ΔCt values across five time points of multiple genes between M and S strains grown under 3µg/ml CYC. T=0 is the time of addition of 1,10-phenanthroline. Time points differ in 15 min with the consecutive one. Each sample represents the mean of three biological replicates, and the error bars represent SD.

**Figure S9**: Comparison of ΔCt values across five time points of multiple genes between M and S strains grown under 20µg/ml FCCP. T=0 is the time of addition of 1,10-phenanthroline. Time points differ in 15 min with the consecutive one. Each sample represents the mean of three biological replicates, and the error bars represent SD.

**Figure S10**: Comparison of ΔCt values across five time points of multiple genes between M-*pbp1Δ* and S-*pbp1Δ* strains grown in YPD and 0.15% H_2_O_2_. T=0 is the time of addition of 1,10-phenanthroline. Time points differ in 15 min with the consecutive one. Each sample represents the mean of three biological replicates, and the error bars represent SD.

**Figure S11**: Comparison of ΔCt values across five time points of multiple genes between M-*puf3Δ* and S-*puf3Δ* strains grown in YPD and 0.15% H_2_O_2_. T=0 is the time of addition of 1,10-phenanthroline. Time points differ in 15 min with the consecutive one. Each sample represents the mean of three biological replicates, and the error bars represent SD.

**Figure S12**: Comparison of ΔCt values across five-time points of multiple genes between M-*pbp1Δpuf3Δ* and S-*pbp1Δpuf3Δ* strains grown in YPD and 0.15% H_2_O_2_. T=0 is the time of addition of 1,10-phenanthroline. Time points differ in 15 min with the consecutive one. Each sample represents the mean of three biological replicates, and the error bars represent SD.

**Figure S13** : Effect of *IVY1*, *MRF1* and *MSD1* on *MKT1* allelic response in CYC, 4NQO and H_2_O_2_. Comparison of relative fitness between *MKT1* alleles in [A-C] *ivy1Δ,* [D-F] *mrf1 Δ* and [G-I] *msd1Δ* backgrounds and the effect of wildtype, *pbp1Δ, puf3Δ* and *pbp1Δpuf3Δ* for each deletion in 0.1µg/ml CYC, 0.3µg/ml 4NQO, and 0.01% H_2_O_2_. The experiments were performed in triplicates, and the error bars represent SD. P-values were calculated using a t-test, and significance was indicated as non-significant (ns), p < 0.05 (*), 0.001 (**), 0.0001 (***) and 0.00001 (****) on the top of each comparison.

**Figure S14**: Allelic phenotype of *MKT1* in Cadmium chloride (CdCl_2_) and High-temperature growth (Htg). 10-fold serial dilution ranging from 10^8^-10^3^ cells/ml of M and S strains were spotted on YPD and the respective growth condition in [A] 300µM CdCl_2_ and [B] incubation at 40°C.

**Figure S15**: Variation of relative fitness difference between M and S backgrounds across the deletion strains. The relative fitness difference (ΔRelative fitness) is the difference between the relative fitness of M and S backgrounds in a particular deletion for each environment. The relative fitness values of wildtype (*Wt*), *cox17Δ, mrs1Δ, rdl2Δ, msd1Δ* , *mrf1Δ* and *ivy1Δ* from the CYC, 4NQO and H_2_O_2_ environments were plotted. The differences were considered for M and S backgrounds, which are statistically significant (p-value < 0.05).

