## Supplementary material for "*MKT1* alleles regulate stress responses through post-transcriptional modulation of Puf3 targets in budding yeast": Figures S1-S15

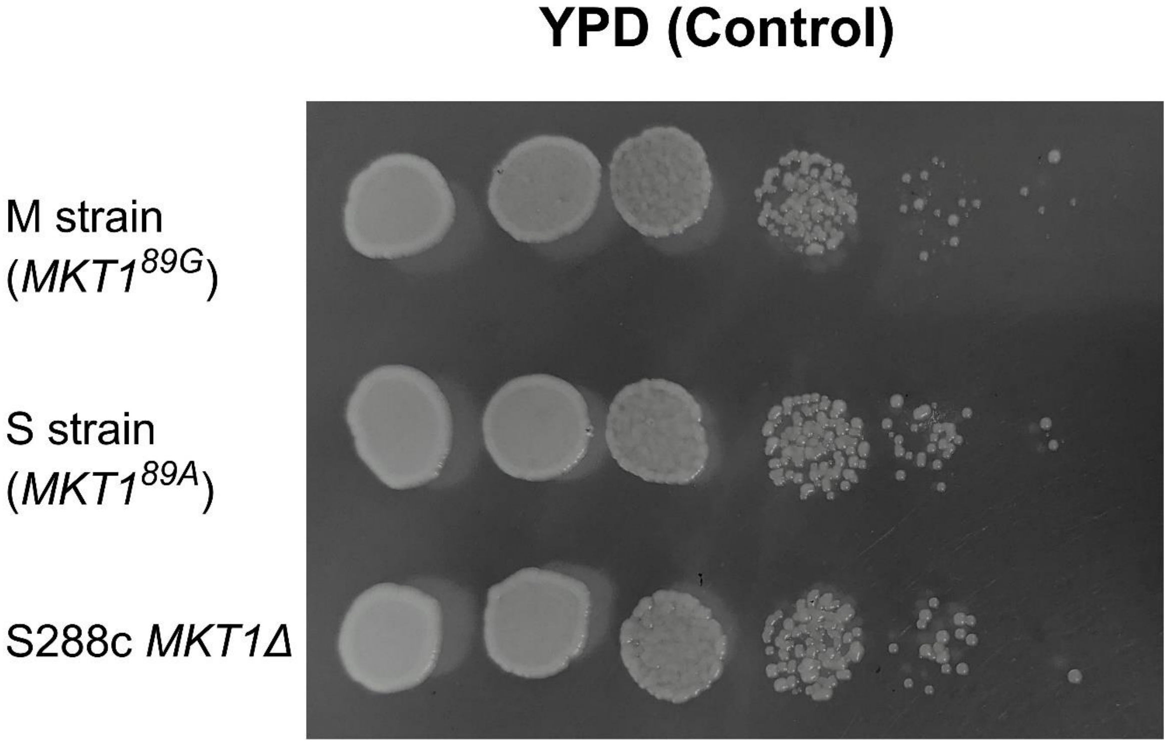


**Figure S1**: Non-essentiality of *MKT1*. 10-fold serial dilution ranging from $10^8$-$10^3$ cells/ml of M, S and S288c *mkt1Δ* strains were spotted on YPD.

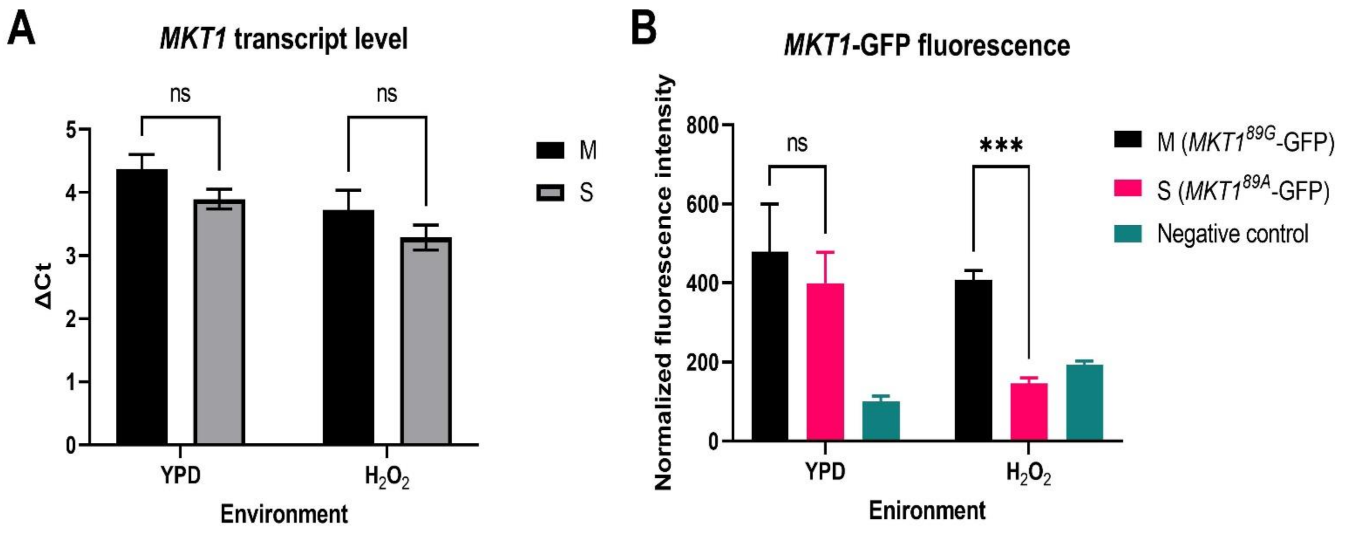


**Figure S2**: Expression of *MKT1* at transcript and protein levels in M and S strains. [A] ΔCt expression levels of *MKT1* in M and S strains grown in YPD control and 0.15% $H_2O_2$. [B] Normalized fluorescence intensity of M (*MKT1$^{89G}$*-GFP), S (*MKT1$^{89A}$*-GFP) and Negative control (S strain) grown in YPD and $H_2O_2$. Strains with GFP-tagged *MKT1* were measured for fluorescence at Em/Ex 485/510 and normalised with their absorbance at 600 nm. Each experiment was performed with three biological replicates, and the resultant mean was plotted, and the error bars represent SD. P-values were calculated using a t-test, and significance was indicated as non-significant (ns), p =< 0.05 (*), 0.001 (**), 0.0001 (***) and 0.00001 (****) on the top of each comparison.

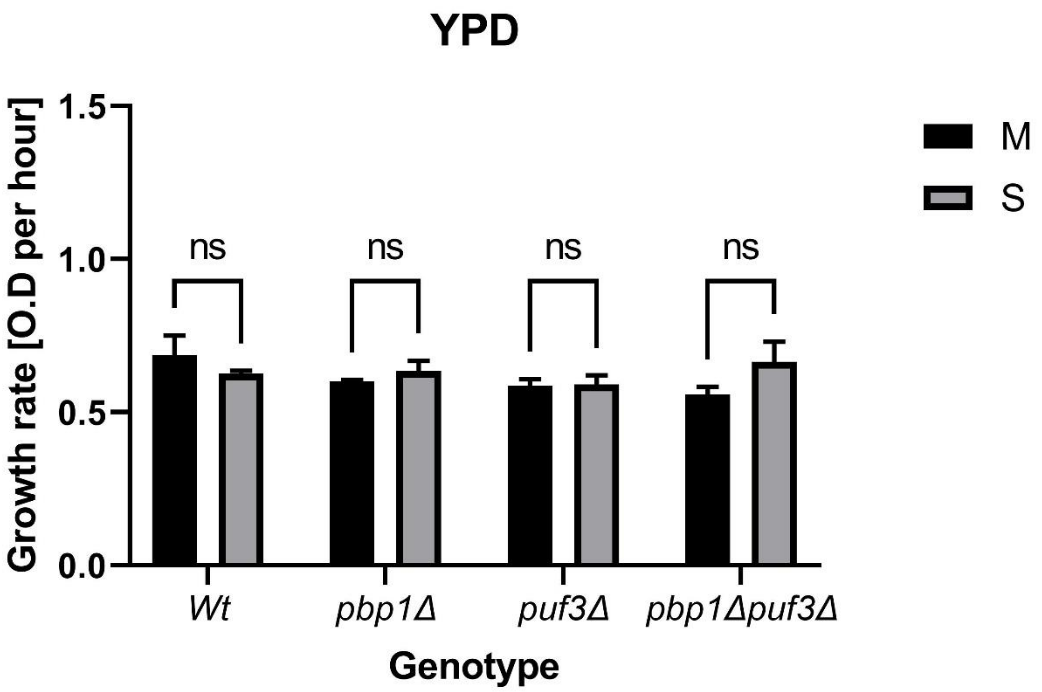


**Figure S3**: Growth rate differences between M and S backgrounds in YPD across *Wt*, *pbp1Δ, puf3Δ* and *pbp1Δpuf3Δ.* The experiments were performed in triplicates, and the error bars represent SD. P-values were calculated using a t-test, and significance was indicated as non-significant (ns) at the top of each comparison.

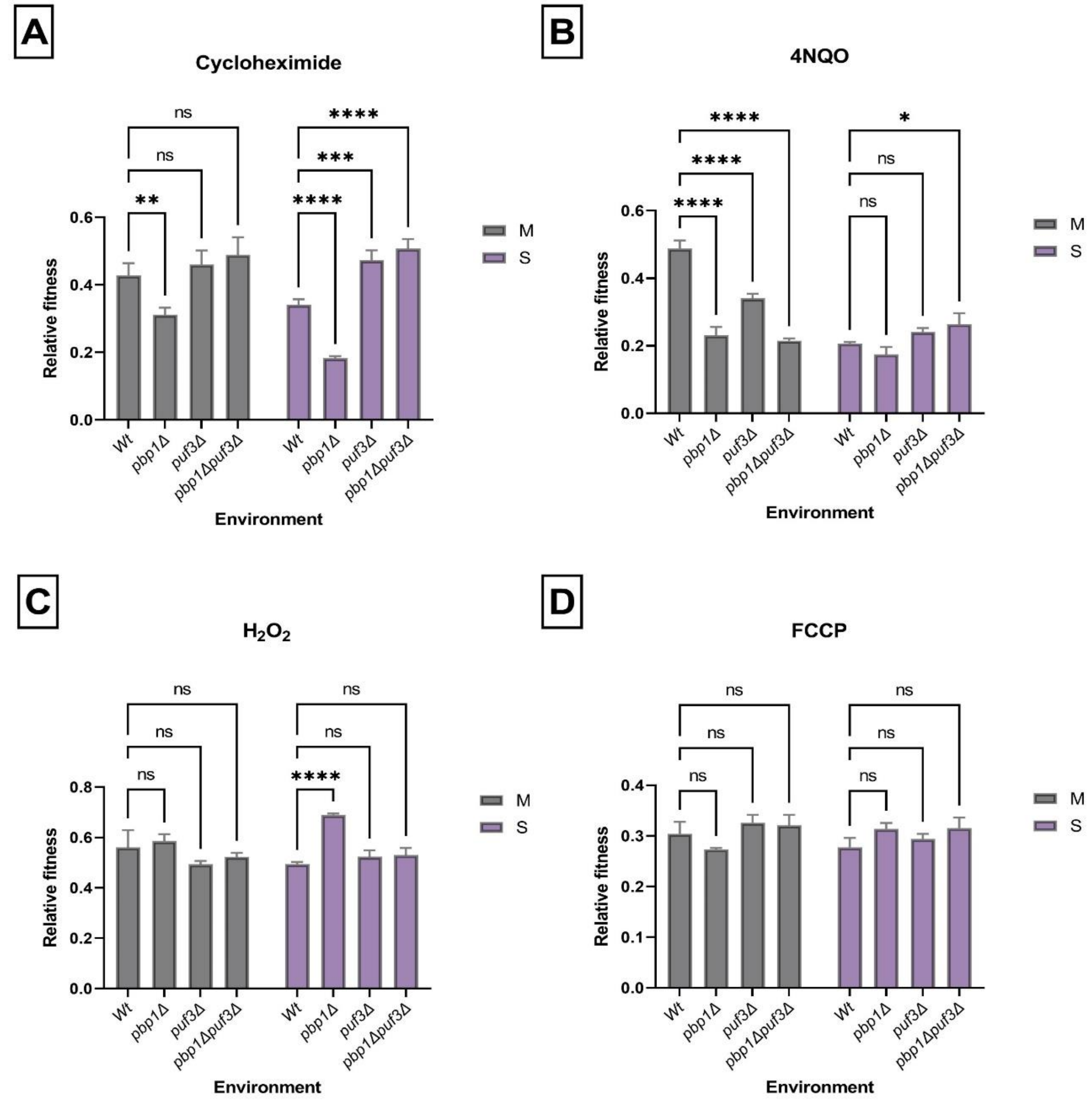


**Figure S4**: Role of *PBP1* and *PUF3* on the fitness of M and S backgrounds in *Wt*, *pbp1Δ, puf3Δ* and *pbp1Δpuf3Δ.* Relative fitness between M and S backgrounds across [A] 0.1 µg/ml CYC, [B] 0.3 µg/ml 4NQO, [C] 0.01% $H_2O_2$ and [D] 1 µg/ml FCCP. The experiments were performed in triplicates, and the error bars represent SD. P-values were calculated using ANOVA, and significance was indicated as non-significant (ns), $p < 0.05$ (*), 0.001 (**), 0.0001 (***) and 0.00001 (****) on the top of each comparison.

#### YPD

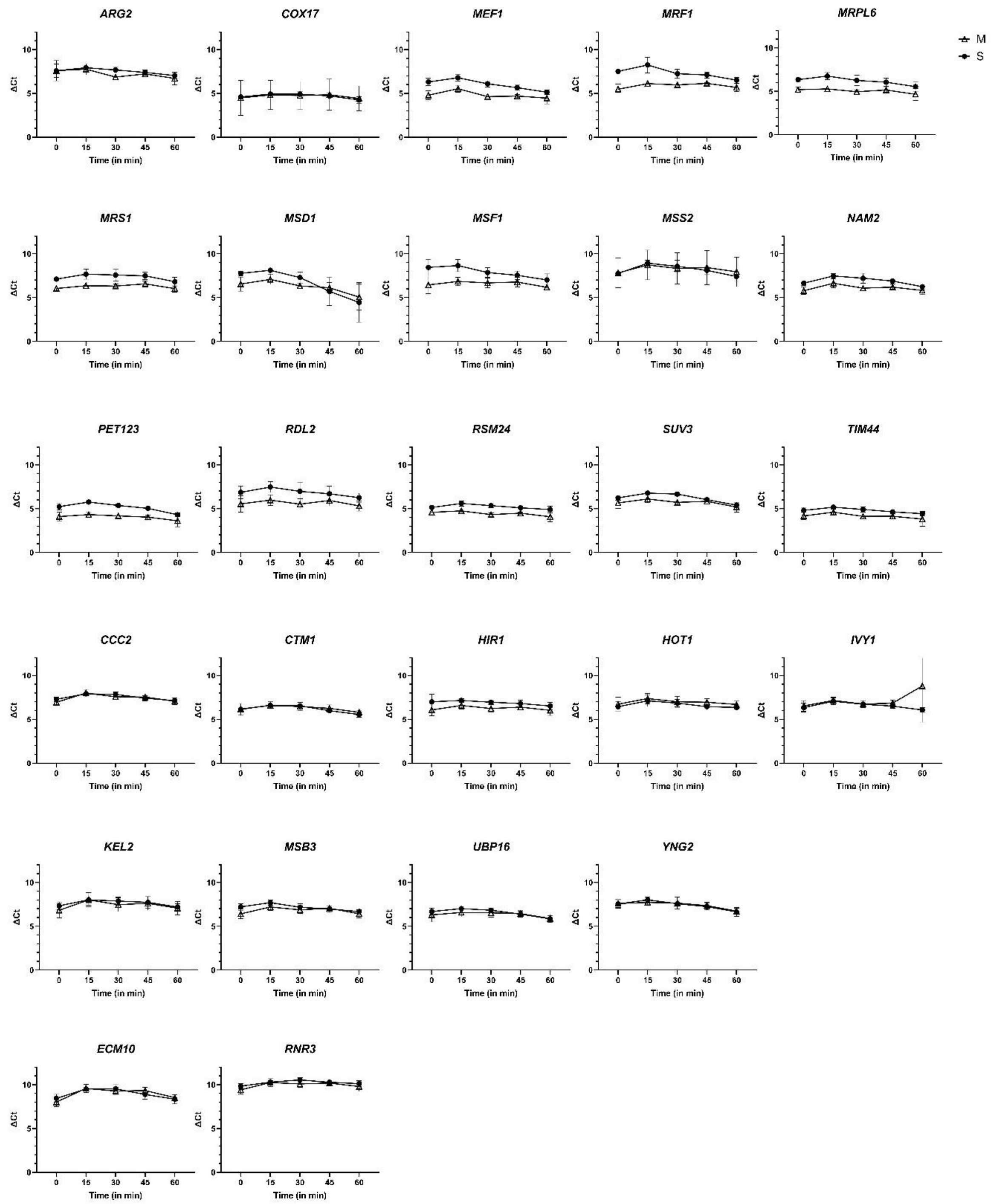


**Figure S5**: Comparison of ΔCt values across five time points of multiple genes between M and S strains grown in YPD. T=0 is the time of addition of 1,10-phenanthroline. Time points differ in 15 min with the consecutive one. Each sample represents as the mean of three biological replicates, and the error bars represent SD.

## $H_2O_2$

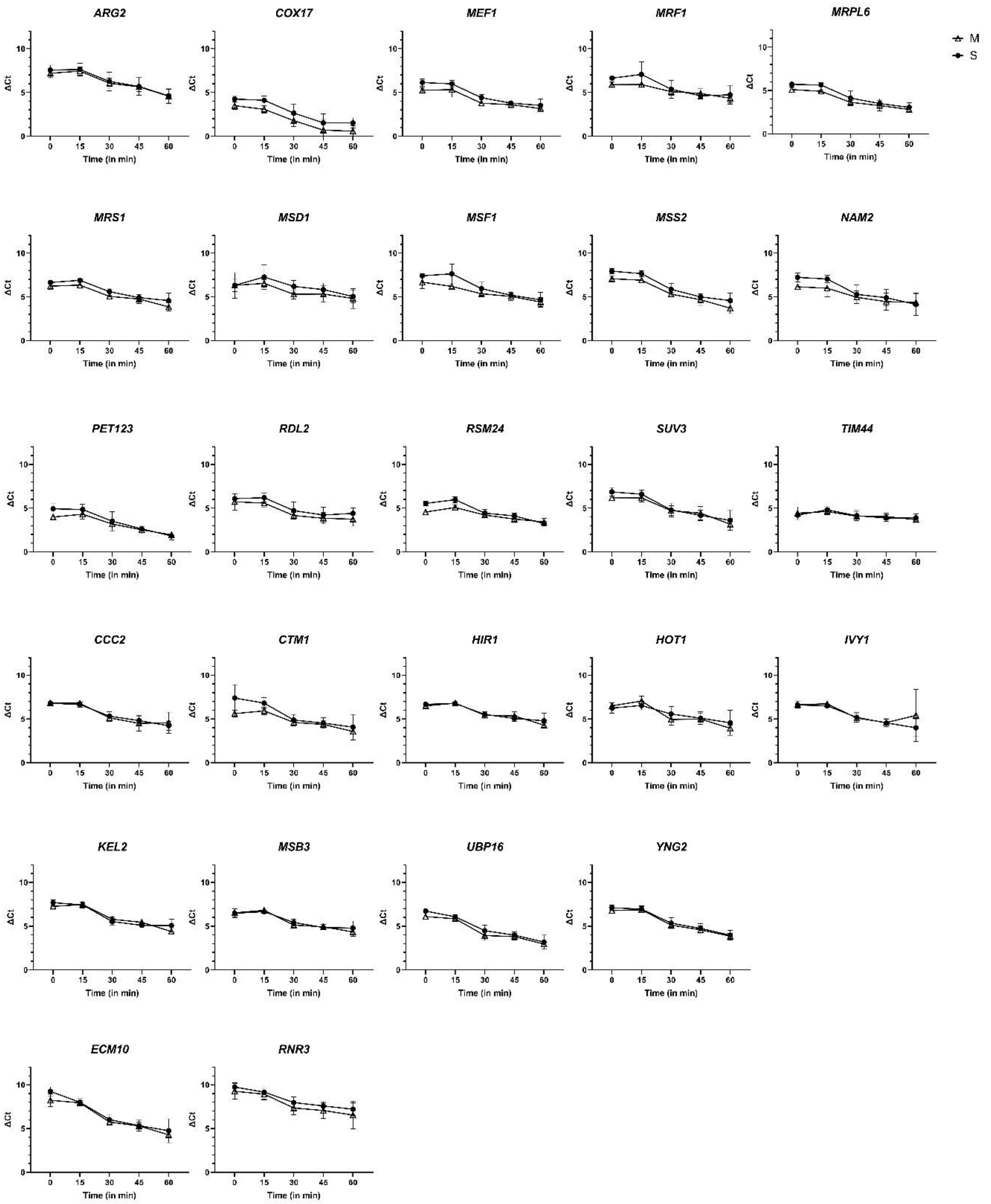


**Figure S6**: Comparison of ΔCt values across five time points of multiple genes between M and S strains grown in 0.15% $H_2O_2$. T=0 is the time of addition of 1,10-phenanthroline. Time points differ in 15 min with the consecutive one. Each sample represents the mean of three biological replicates, and the error bars represent SD.

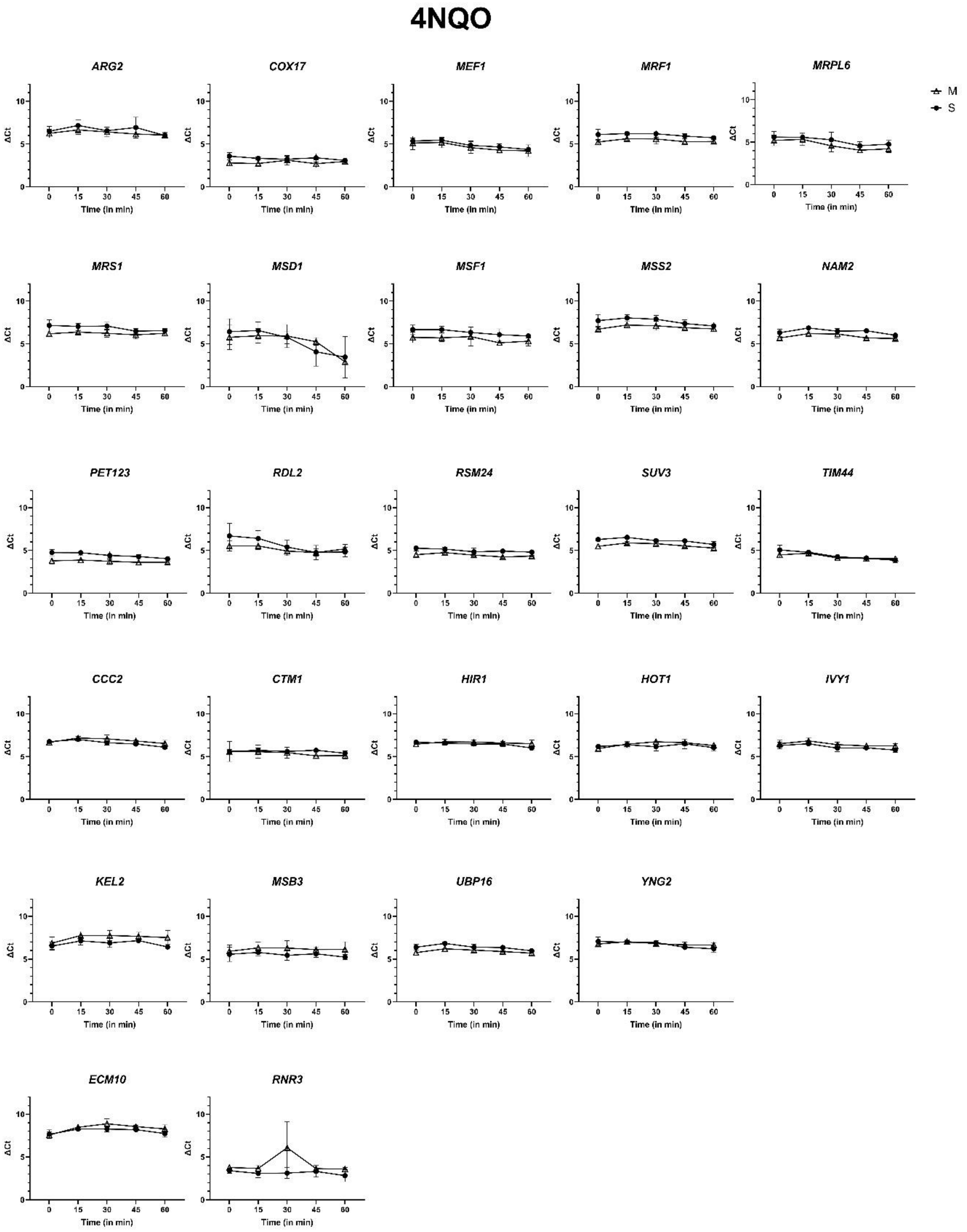


**Figure S7**: Comparison of ΔCt values across five time points of multiple genes between M and S strains grown under 2µg/ml 4NQO. T=0 is the time of addition of 1,10-phenanthroline. Time points differ in 15 min with the consecutive one. Each sample represents the mean of three biological replicates, and the error bars represent SD.

### Cycloheximide

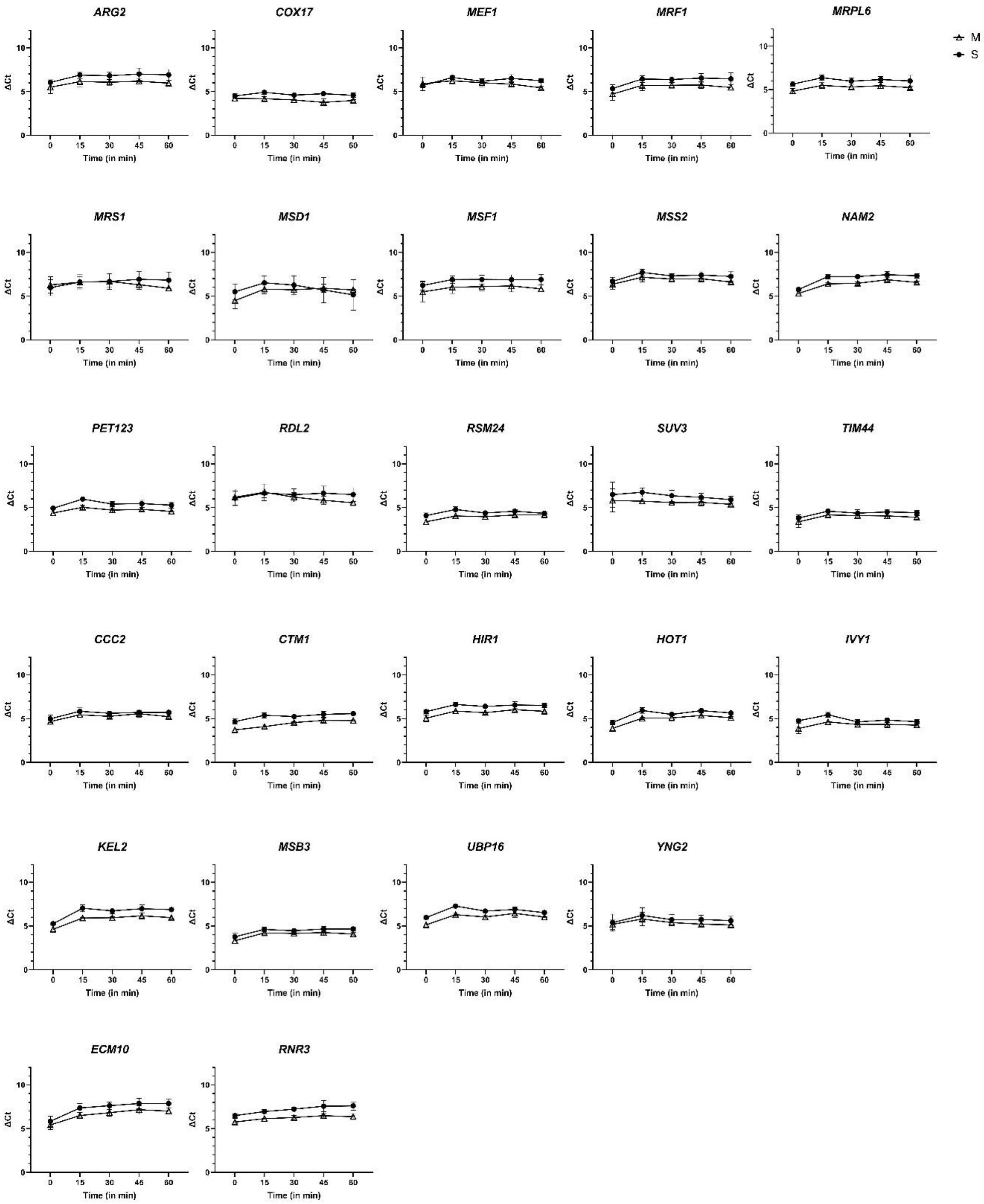


**Figure S8**: Comparison of ΔCt values across five time points of multiple genes between M and S strains grown under 3μg/ml CYC. T=0 is the time of addition of 1,10-phenanthroline. Time points differ in 15 min with the consecutive one. Each sample represents the mean of three biological replicates, and the error bars represent SD.

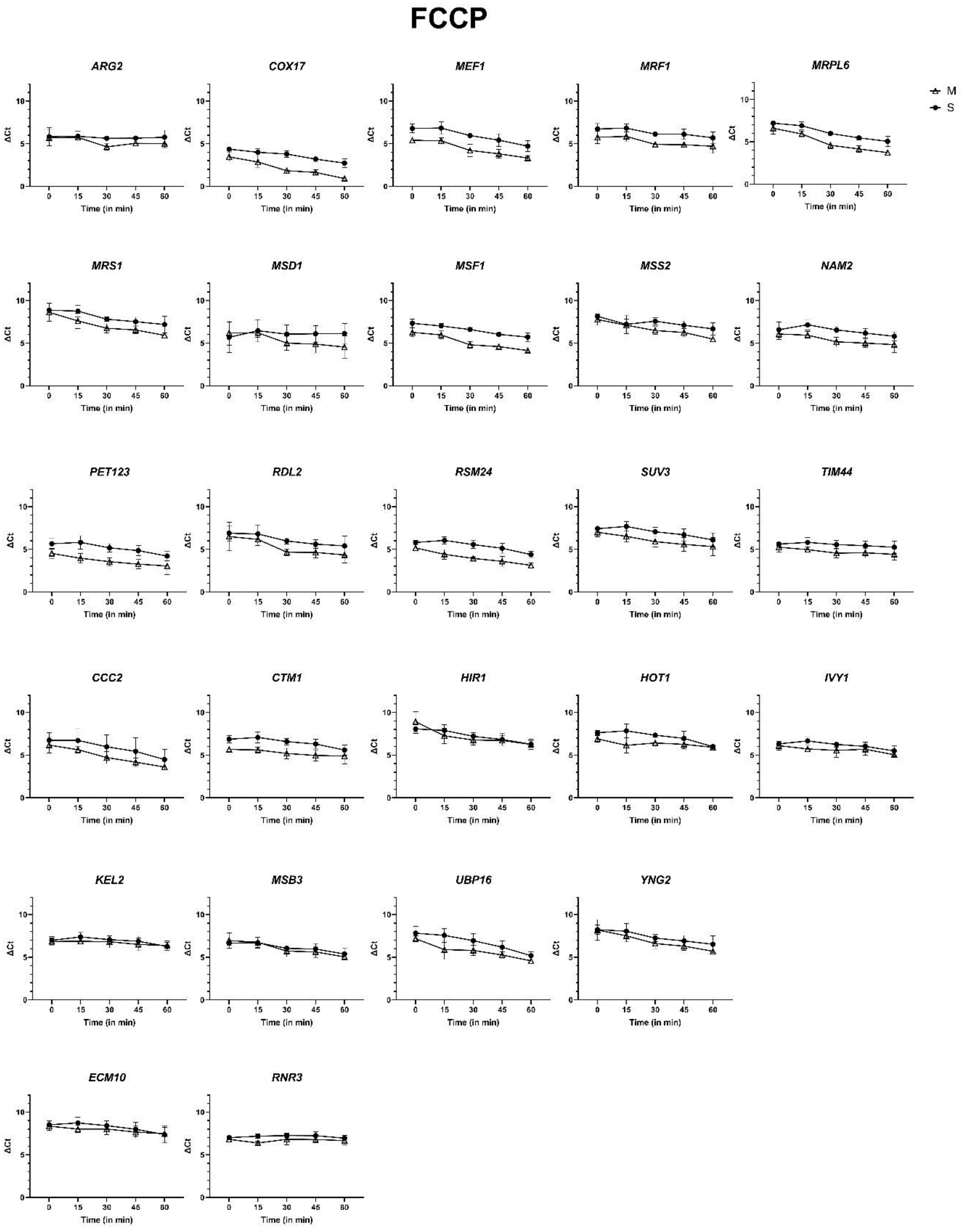


**Figure S9**: Comparison of ΔCt values across five time points of multiple genes between M and S strains grown under 20µg/ml FCCP. T=0 is the time of addition of 1,10-phenanthroline. Time points differ in 15 min with the consecutive one. Each sample represents the mean of three biological replicates, and the error bars represent SD.

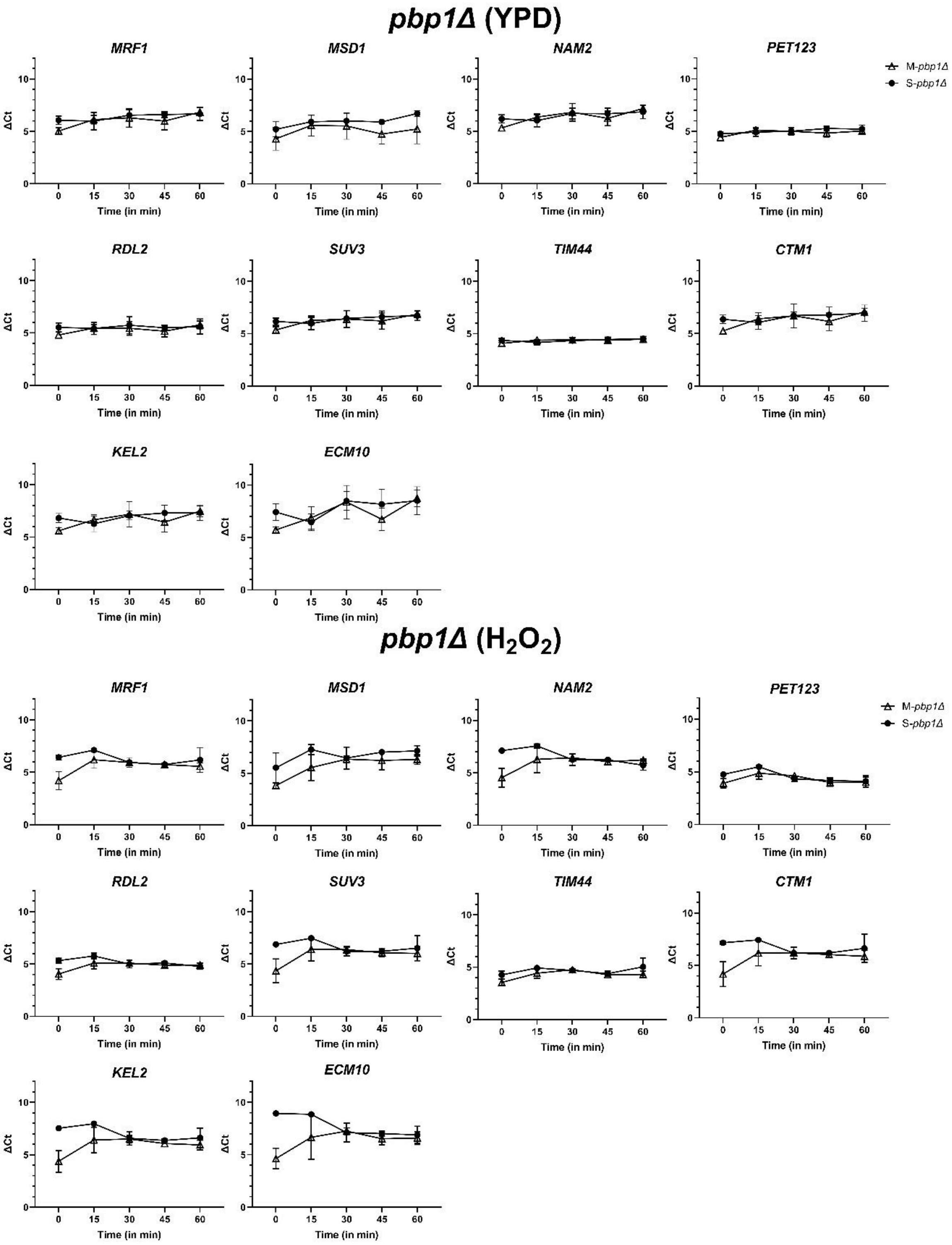


**Figure S10**: Comparison of ΔCt values across five time points of multiple genes between M-*pbp1Δ* and S-*pbp1Δ* strains grown in YPD and 0.15% $H_2O_2$. T=0 is the time of addition of 1,10-phenanthroline. Time points differ in 15 min with the consecutive one. Each sample represents the mean of three biological replicates, and the error bars represent SD.

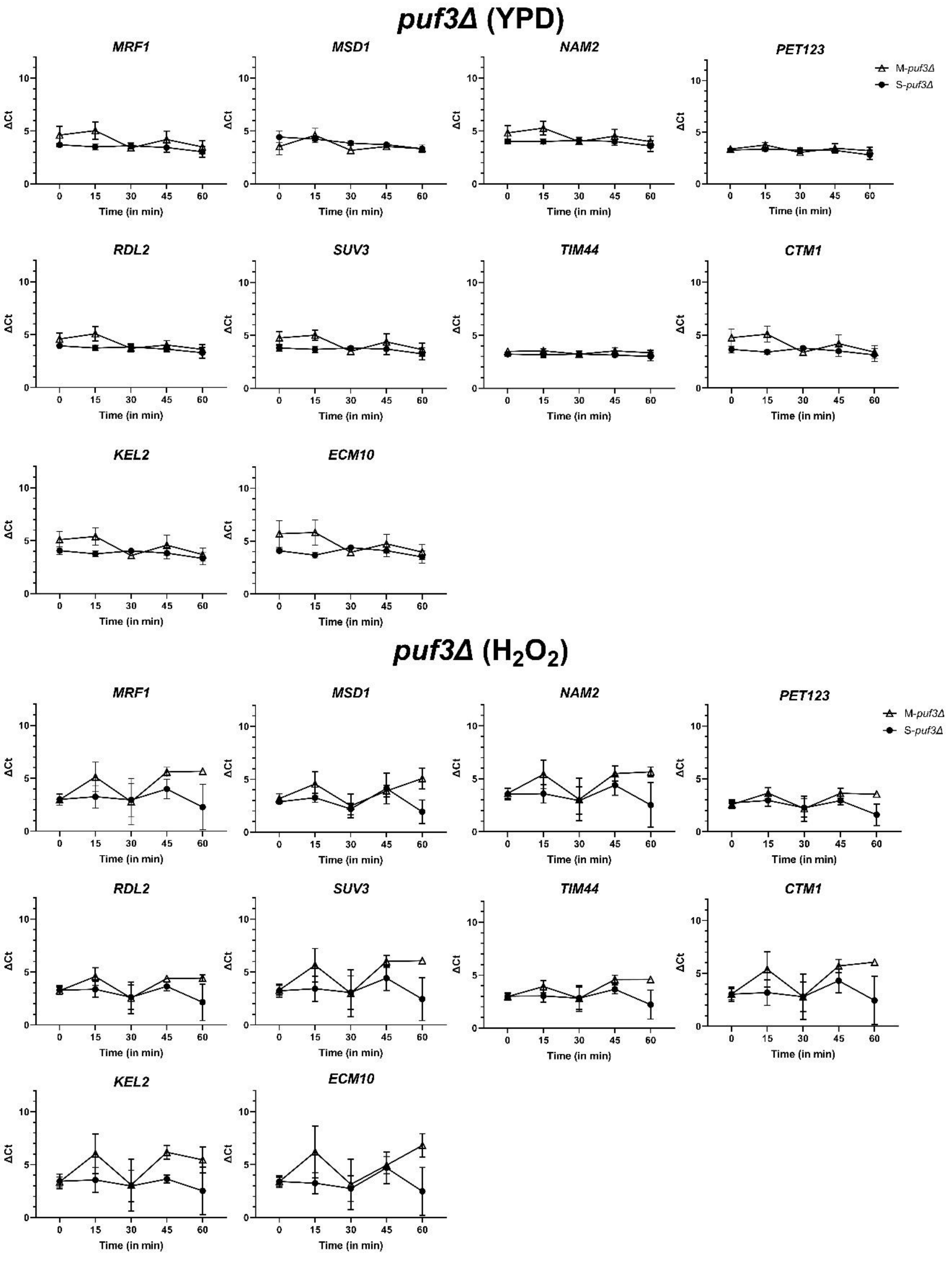


**Figure S11**: Comparison of ΔCt values across five time points of multiple genes between M-*puf3Δ* and S-*puf3Δ* strains grown in YPD and 0.15% $H_2O_2$. T=0 is the time of addition of 1,10-phenanthroline. Time points differ in 15 min with the consecutive one. Each sample represents the mean of three biological replicates, and the error bars represent SD.

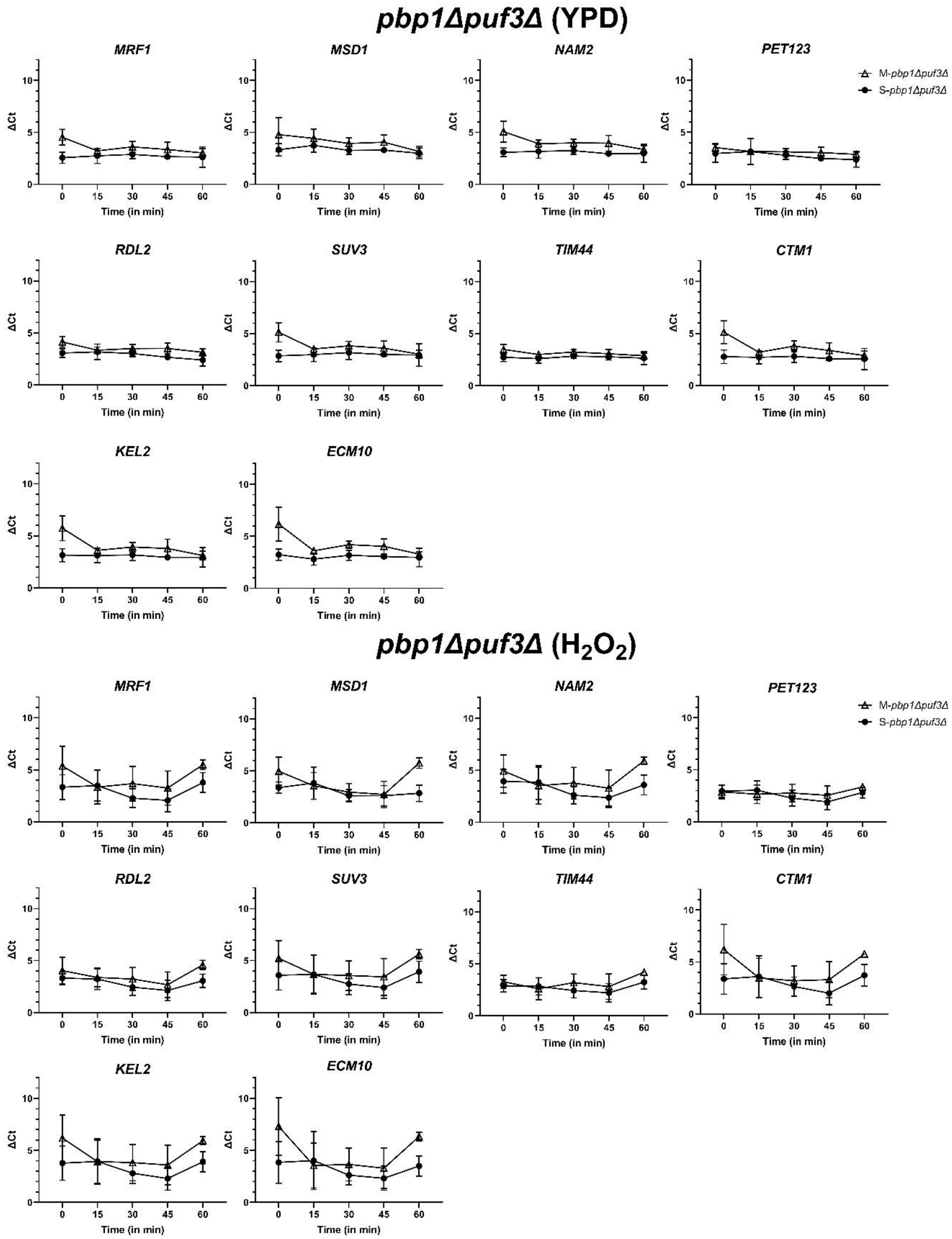


**Figure S12**: Comparison of ΔCt values across five-time points of multiple genes between M-*pbp1Δpuf3Δ* and S-*pbp1Δpuf3Δ* strains grown in YPD and 0.15% $H_2O_2$. T=0 is the time of addition of 1,10-phenanthroline. Time points differ in 15 min with the consecutive one. Each sample represents the mean of three biological replicates, and the error bars represent SD.

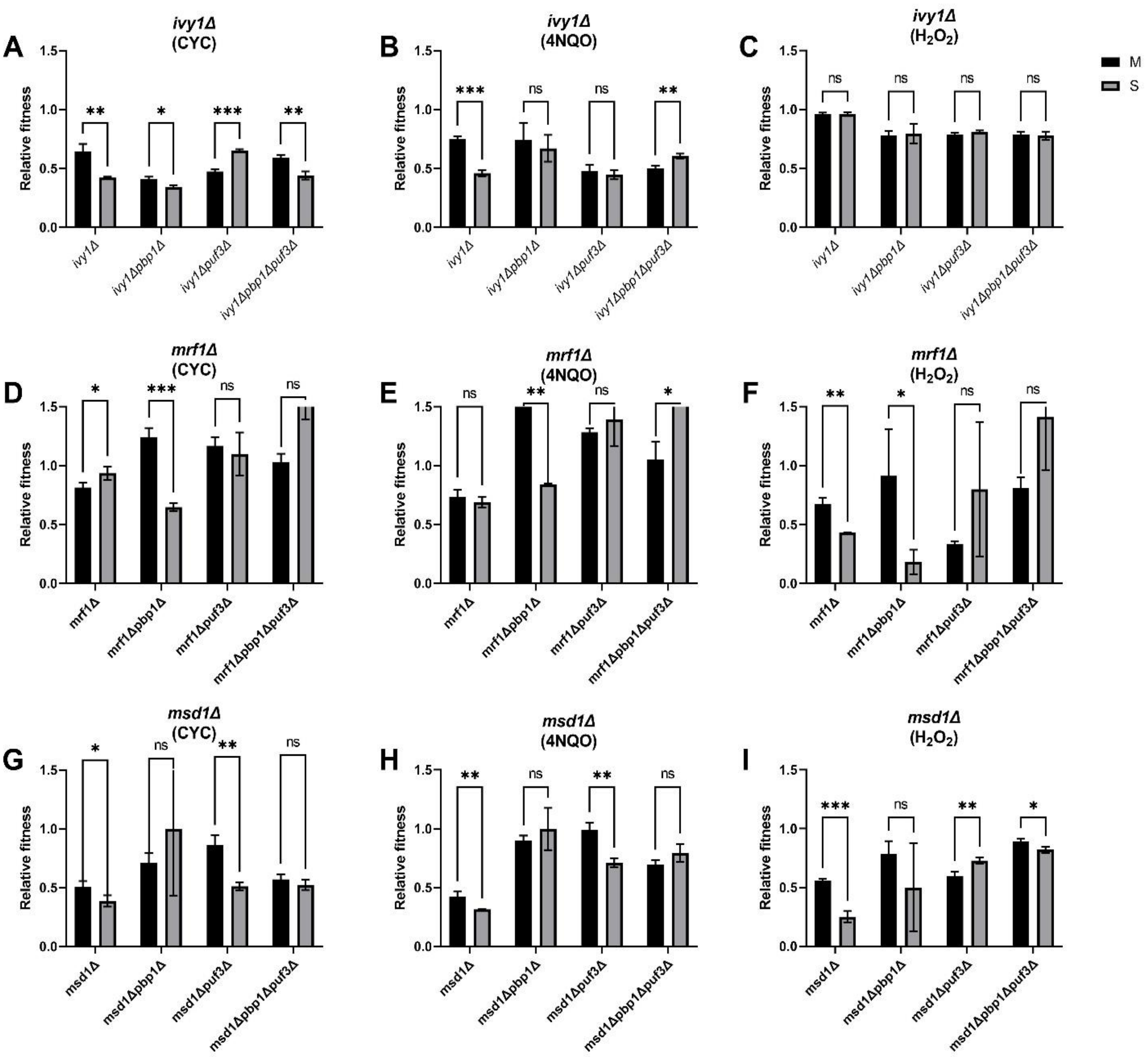


**Figure S13**: Effect of *IVY1*, *MRF1* and *MSD1* on *MKT1* allelic response in CYC, 4NQO and $H_2O_2$. Comparison of relative fitness between *MKT1* alleles in [A-C] *ivy1Δ,* [D-F] *mrf1Δ* and [G-I] *msd1Δ* backgrounds and the effect of wildtype, *pbp1Δ, puf3Δ* and *pbp1Δpuf3Δ* for each deletion in 0.1µg/ml CYC, 0.3µg/ml 4NQO, and 0.01% $H_2O_2$. The experiments were performed in triplicates, and the error bars represent SD. P-values were calculated using a t-test, and significance was indicated as non-significant (ns), $p < 0.05$ (*), 0.001 (**), 0.0001 (***) and 0.00001 (****) on the top of each comparison.

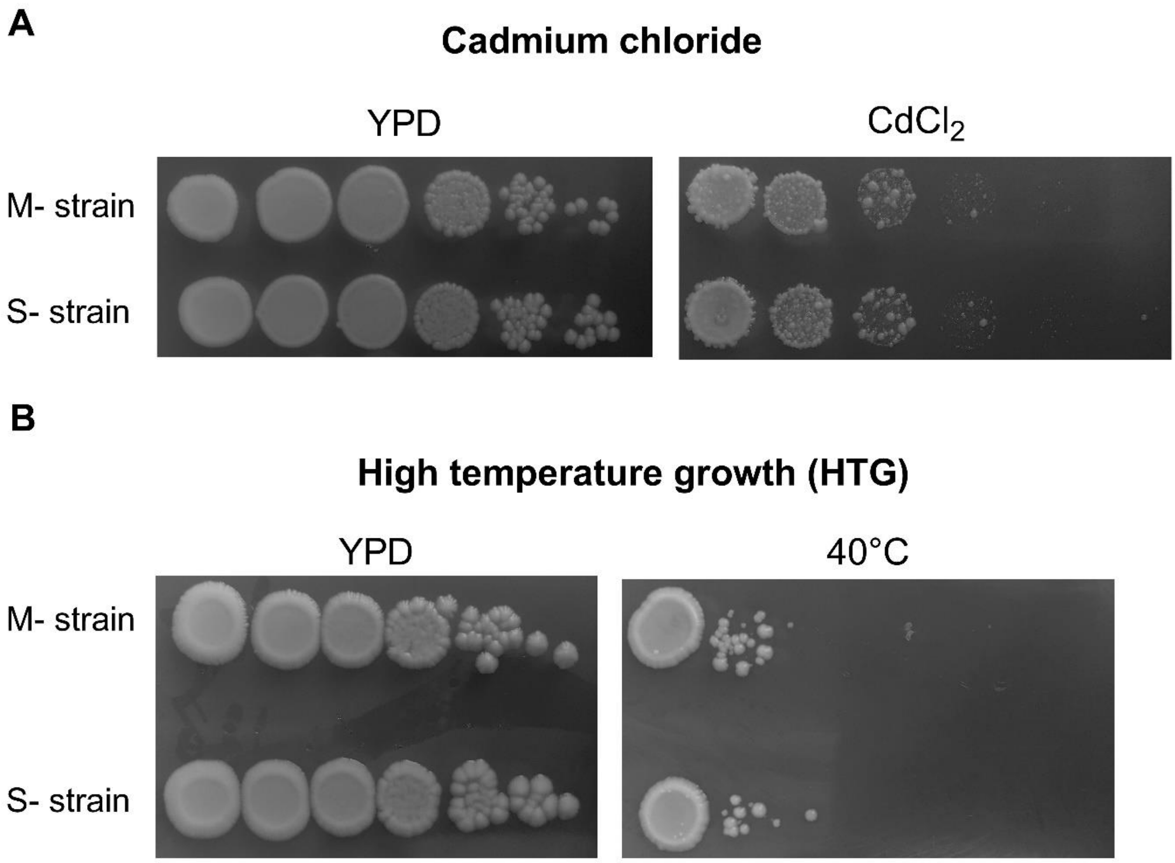


**Figure S14**: Allelic phenotype of *MKT1* in Cadmium chloride ($CdCl_2$) and High-temperature growth (Htg). 10-fold serial dilution ranging from $10^8$-$10^3$ cells/ml of M and S strains were spotted on YPD and the respective growth condition in [A] 300µM $CdCl_2$ and [B] incubation at 40°C.

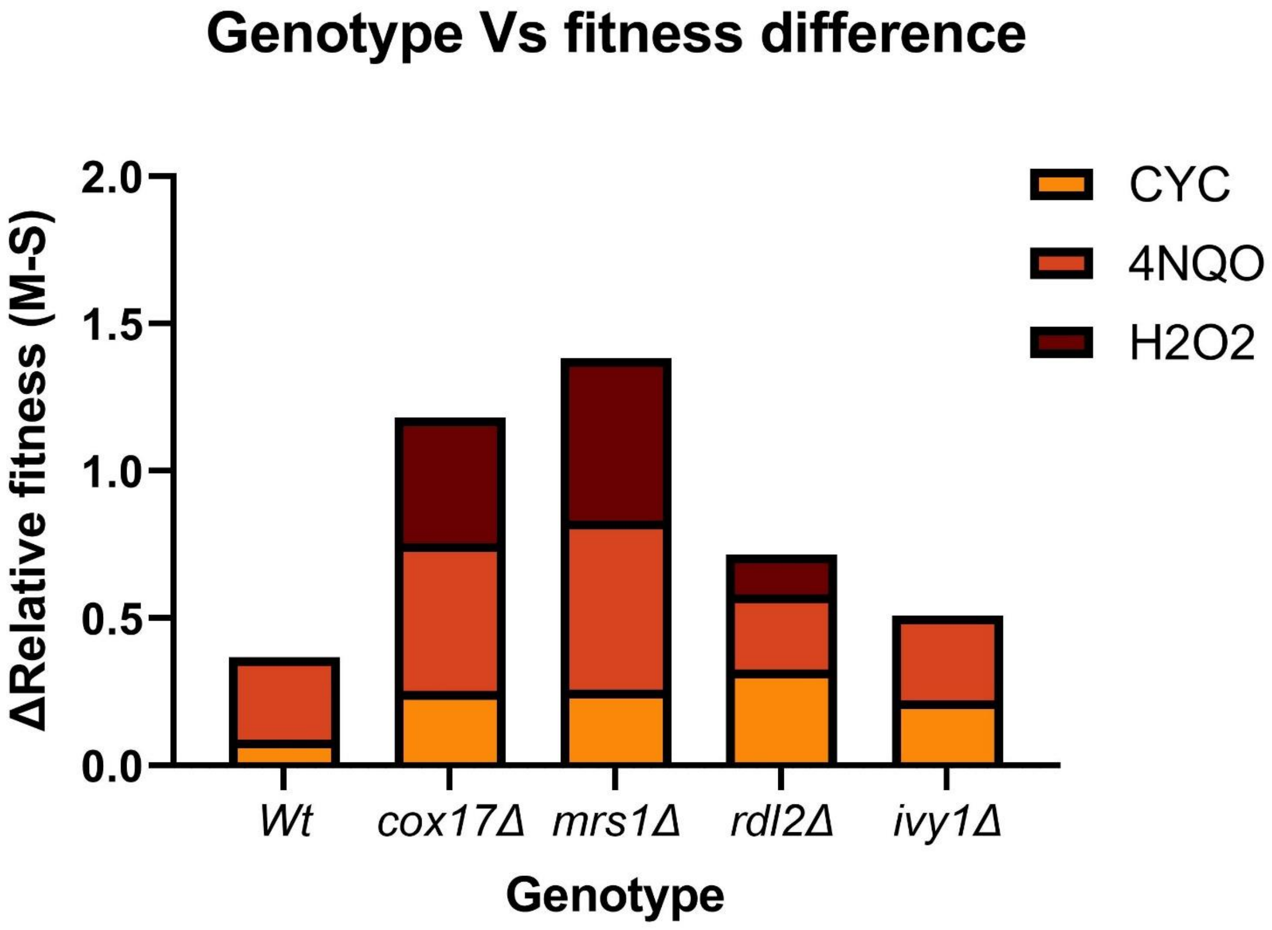


**Figure S15**: Variation of relative fitness difference between M and S backgrounds across the deletion strains. The relative fitness difference (ΔRelative fitness) is the difference between the relative fitness of M and S backgrounds in a particular deletion for each environment. The relative fitness values of wildtype (*Wt*), *cox17Δ, mrs1Δ, rdl2Δ, msd1Δ*, *mrf1Δ* and *ivy1Δ* from the CYC, 4NQO and $H_2O_2$ environments were plotted. The differences were considered for M and S backgrounds, which are statistically significant ($p < 0.05$).
